# Cancer drug discovery as a low rank tensor completion problem

**DOI:** 10.1101/2021.03.08.434311

**Authors:** Vasanth S. Murali, Didem Ağaç Çobanoğlu, Michael Hsieh, Meyer Zinn, Venkat S. Malladi, Jonathan Gesell, Noelle S. Williams, Erik S. Welf, Ganesh V. Raj, Murat Can Çobanoğlu

## Abstract

The heterogeneity of cancer necessitates developing a multitude of targeted therapies. We propose the view that cancer drug discovery is a low rank tensor completion problem. We implement this vision by using heterogeneous public data to construct a tensor of drug-target-disease associations. We show the validity of this approach computationally by simulations, and experimentally by testing drug candidates. Specifically, we show that a novel drug candidate, SU11652, controls melanoma tumor growth, including BRAF^WT^ melanoma. Independently, we show that another molecule, TC-E 5008, controls tumor proliferation on *ex vivo* ER+ human breast cancer. Most importantly, we identify these chemicals with only a few computationally selected experiments as opposed to brute-force screens. The efficiency of our approach enables use of *ex vivo* human tumor assays as a primary screening tool. We provide a web server, the Cancer Vulnerability Explorer (accessible at https://cavu.biohpc.swmed.edu), to facilitate the use of our methodology.

## 1. Introduction

Cancer is characterized by genomic variation (Hanahan and Weinberg, 2011) that leads to intra- (Meacham and Morrison, 2013) and inter-tumoral heterogeneity (Holohan et al., 2013). Therefore cancer drug discovery is stratified by cancer type and, recently, subtype (Dugger et al., 2017). Hence there is a need for a wide array of cancer drug discovery projects that pursue different subtypes of the disease.

Recently, however, there has emerged the realization that drug discovery productivity has declined over the decades (Paul et al., 2010), and exponentially so (Scannell et al., 2012). The cause of this decline has been attributed to a drop in the clinical predictive value of reductionist screening practices (Scannell and Bosley, 2016). While multiple clinically relevant functional assays exist (Friedman et al., 2015), these can achieve a throughput of only 10^1^ to 10^2^, which is orders of magnitude lower than necessary for brute-force screening: 10^5^ to 10^6^(Macarron et al., 2011).

The result of these two trends creates a two-pronged dilemma: we need more drug discovery projects, while using functional assays with lower throughput. The solution we propose is to computationally select a small number of exceptionally well-suited drug candidates. These few drugs can then be tested in low throughput assays with high clinical relevance (3D or *ex vivo*). Any hits that emerge from the proposed mechanism would, by virtue of the assays, have a high likelihood of clinical translation. However the key challenge remains identification of the promising drug candidates.

We hereby present the argument that synthesizing diverse types of public data into a tensor of drug-target-disease associations can efficiently identify targeted drug candidates. There are several advantages to this representation. Mathematically, representing the multitude of diseases that comprise cancer in this manner allows for information sharing across related cancer types (eg: adenocarcinomas of different tissues). Conceptually, it serves as a unification of currently disparate datasets that are in actuality related to each other for cancer drug discovery purposes.

The practice of constructing a drug-target-disease tensor relies heavily on the definition of “disease.” In this study we adopt two separate definitions: tissue lineage (Visvader, 2011) and mutation (Lawrence et al., 2013). These are not exclusive; instead, a disease can be defined as a composition of the two (ex: BRAF^V600E^ melanoma). While there are nuances to each of these definitions, as well as other possible ways of defining cancer subtypes, we consider that these two definitions cover most common ways of defining cancer in research as well as in the clinic.

The linkage of diseases to protein targets of significance is another critical step. To this end, we benefit from a number of recent perturbation studies such as DRIVE (McDonald et al., 2017) or Achilles (Tsherniak et al., 2017). In previous datasets, such as gene expression or protein abundance measurements, the abundance of a gene or protein is measured as a proxy. However those datasets provide no knowledge about any compensatory mechanisms or essentiality, either of which would render a gene product unsuitable as a drug target. In contrast, DRIVE and Achilles differ significantly from previous datasets in that they measure the response of a relatively high number of cell lines to genetic perturbations. By measuring the outcome of a perturbation, these latest generation of datasets allow us to search for genes whose loss of function is specifically lethal to a targeted subtype of cancer yet not lethal for a wide array of other cell lines. We present a statistical method, based on hypergeometric enrichment testing, to extract such targeted disease-target links from the perturbation datasets.

Finally, to turn these pathobiological hypotheses to clinically actionable drug candidates, we leverage large scale chemical-protein association data. We use the STITCH (Szklarczyk et al., 2015) dataset for this purpose since it “stitches” together 14 different databases that serve this type of data. We use Ensembl (Hubbard, 2002) to link the proteins that STITCH reports to the genes that are perturbed in the datasets such as DRIVE. We finally assess the druglikeness of the chemicals by calculating their chemical similarity to approved drugs in DrugBank (Wishart et al., 2017). In the end, we create a tripartite graph of drug-target-disease links, which can mathematically be represented as a third-order tensor.

We can leverage the tensor representation to make a large number of useful predictions through low rank tensor completion. A drug-target-disease tensor constructed with Cancer Cell Line Encyclopedia (CCLE) (Barretina et al., 2012) subtype annotations as “diseases,” and some basic drug-likeness filters yields 13,190 associations among 8,211 chemicals, 523 proteins, and 21 diseases. This tensor is extremely sparse: less than 0.015% of its possible entries are known. Knowing this tensor in full is highly desirable since it would allow us to answer multiple types of queries such as potential new drug candidates, or novel mechanisms of action.

We solve this challenge through low rank tensor completion. Briefly, we decompose this tensor as a sum of outer products (Hitchcock, 1927), a method also known as CANDECOMP/PARAFAC (CP) decomposition (Long et al., 2019). Low rank tensor completion via factorization, while novel to the cancer domain, has been proposed to fill similar extremely sparse tensors for large consumer-facing technology companies such as Netflix (users-movies-time, ratings tensor) a decade ago (Xiong et al., 2010). Likewise, both we (Cobanoglu et al., 2013) and others (Gonen and Kaski, 2014, Liu et al., 2016, Hao et al., 2017) have previously used matrix factorization to complete very sparse drug-target interaction matrices. In light of this literature, our work is an extension from completing drug-target interaction matrices to drug-target-disease tensors. The mathematical fundamentals in all such work is the decomposition of a given set of associations organized as a multidimensional array (matrix or tensor) into subcomponents such that their multiplication then returns the original object with high fidelity. The inferred subcomponents are latent variables that enable the prediction of any *a priori* unknown entry in the original tensor. In this manner, we can complete the targeted cancer drug discovery tensor despite a high degree of sparsity.

We validated our approach by finding drug candidates against melanoma and ER+ breast cancer. To find the melanoma drug candidate, SU11652, we used a 3D *in vitro* assay that we had previously developed (Murali et al., 2019). We then validated our prediction in multiple *in vivo* models, including BRAF^WT^ melanoma which is clinically challenging. To find the ER+ breast cancer drug candidate, we used *ex vivo* human tumor culture, which is arguably the golden standard assay in terms of clinical relevance. We successfully found drug candidates with these low throughput assays only due to our computational lead selection. To render our work more accessible, we also built a web server that we call the Cancer Vulnerability Explorer (or CaVu in short) (http://cavu.biohpc.swmed.edu), so that any researcher can find drug or target candidates for their cancer subtype of interest using our methodology.

## 2. Results

### 2.1. Defining cancer subtypes

Cancer is a disease caused and characterized by genomic lesions, therefore mutations are commonly used to define cancer subtypes (Dey et al., 2017). In the clinic, mutations are used as biomarkers for targeted therapies, such as in the use of vemurafenib against melanoma harboring the BRAF^V600E^ mutation (Li et al., 2013, Chapman et al., 2011). Mutations are also used to define subclones within a single patient in order to explain resistance (Schmitt et al., 2015). Some somatic mutations can also be cancer drivers, with 568 genes currently known to have the potential to adopt such a role (Martinez-Jimenez et al., 2020). In conclusion, mutations are fundamental to our understanding and categorization of cancer.

Tissue lineage of cancer is also commonly used to classify cancer types and subtypes since lineage influences the tumor behavior and response to therapy (Visvader, 2011, Garraway and Sellers, 2006). The guidance of tissue lineage on cancer drug discovery started with the NCI-60 program (Chabner, 2016), and has led to more recent efforts such as the Genomics of Drug Sensitivity in Cancer (GDSC) (Garnett et al., 2012, Iorio et al., 2016), the CCLE (Barretina et al., 2012), and the Cancer Therapeutics Response Portal (CTRP) (Basu et al., 2013, Seashore-Ludlow et al., 2015). The tissue lineage of cancer has a long history in shaping the search for potential therapeutic agents.

In our web server, we use both of these definitions of subtypes, i.e. mutations and histology, to allow users to specify any cancer subtype of interest to them. If the user chooses to define subtypes based on histology, then CaVu shows a list of the histological type and subtype annotations, tumor site annotations, pathologist annotations and other relevant information for the list of cell lines available for vulnerability analysis based on CCLE (Barretina et al., 2012) data (“cell line annotations” v20181226, latest version as of the writing). Using this functionality, the users can pose queries such as “lung adenocarcinomas” or “all adenocarcinomas”. When making our melanoma drug predictions, we relied on the tissue lineage based subtype definition.

Alternatively, in CaVu we also offer the option to define subtypes on the basis of specific mutations. When choosing this option, the users start by picking their gene of interest. We then display a list of all the known mutations in that gene in all the cell lines in our integrated analysis platform. To do this, we use the CCLE merged mutation calls (v20180502, which is the latest version as of the writing). The user then optionally filters for cell lines that fit their definition of a cancer subtype, or simply uses the automated annotation (Figure 1A). Most cell lines have relatively few mutations, making subtype definitions more likely to be precise (Figure 1B). We then use this collection of cell lines as the basis for our vulnerability analysis. When making the ER+ breast cancer drug predictions, we used the genomic annotations of cell lines to categorize the breast cancer cell lines.

**Figure 1:**
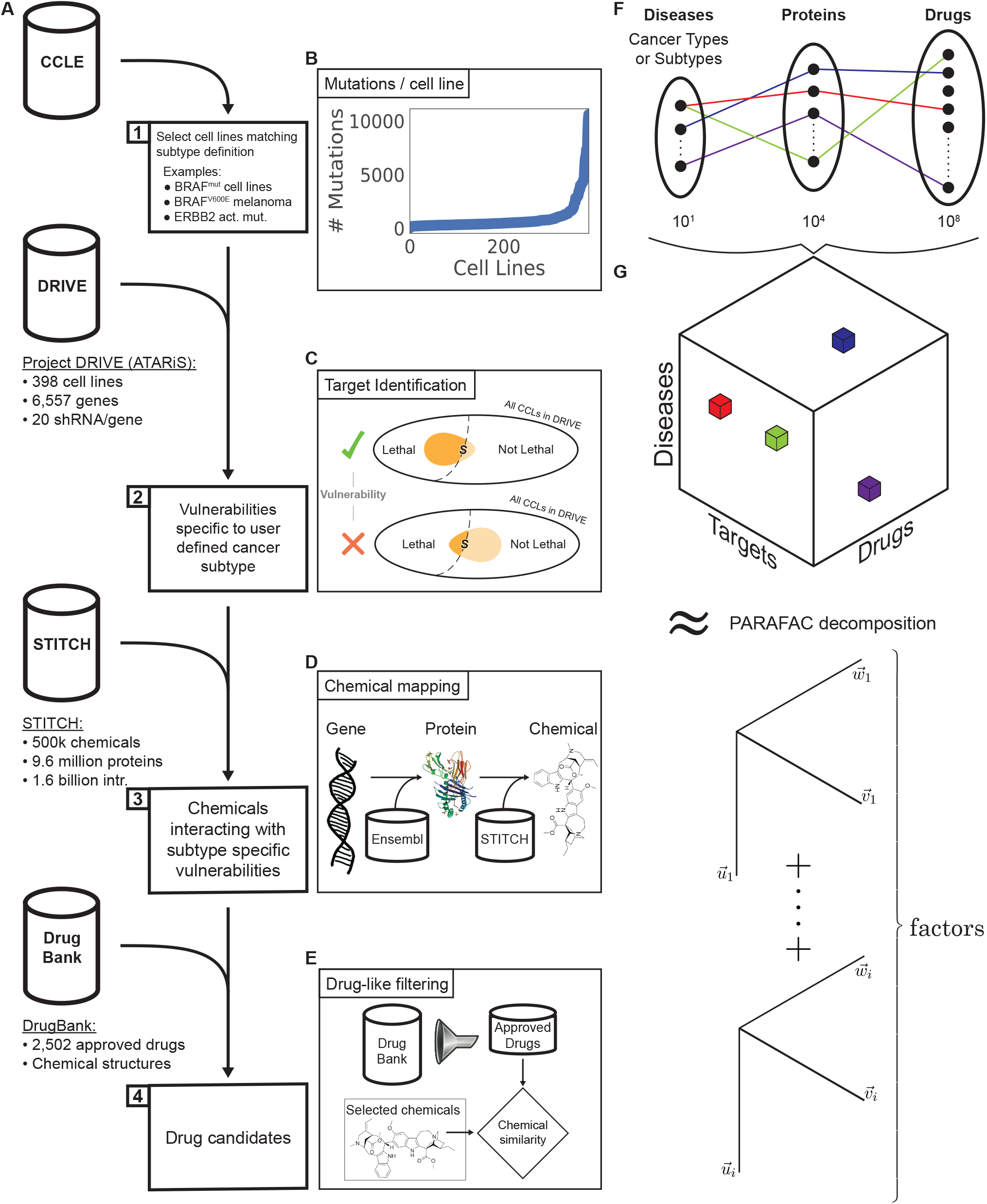
CaVu schematic overview. When using CaVu, users first define a cancer subtype of interest based on mutations (ex: BRAF^V600E^ mutated) and/or lineage (data from CCLE). Then, we automatically identify the subtype specific vulnerabilities using Project DRIVE data by running a statistical test. Next, we link the significant vulnerabilities to chemicals (STITCH data) and filter for drug-like molecules (DrugBank data). To predict the unknown but most likely drug-target-disease associations, we then use low rank tensor completion on a drug-target-disease interaction tensor (using PARAFAC decomposition). These predictions yield multiple benefits; our main objective being the identification of unknown yet likely targets to explain mechanism of action.

### 2.2. Identifying vulnerabilities for a cancer subtype

Given a cancer subtype represented by a set of cell lines, we identify oncogene addictions (i.e. targets) using the hypergeometric enrichment test (Figure 1C). Briefly, this test checks whether a knockdown is lethal to members of the predefined set more often than one would expect by chance. Since we test thousands of hypotheses at once, we use the Benjamini-Hochberg method to control the false discovery rate (FDR) (Benjamini and Hochberg, 1995). In our interactive web interface, we present both the uncorrected *p*-value, as well as FDR, which we call the *q*-value.

Our approach searches for genes whose knockdown is particularly lethal to the cell lines in the user defined subtype, as opposed to all the available cancer cell lines. The knockdown of any non-specific target will be lethal to many cell lines, including those outside the user-defined subtype. Therefore they will not be enriched and score poorly. We use the cell lines that are not in the target subtype, collectively, as a filter to eliminate non-specific targets. Our analysis prioritizes vulnerabilities specific to a particular category of cell lines over generally toxic knockdown targets, such as essential housekeeping genes. Thus our tool enables the user to rapidly identify the most interesting targets for their cancer subtype of interest.

The synthetic lethality data is critical for our vulnerability identification pipeline. There are currently two large-scale synthetic lethality screens: DRIVE (McDonald et al., 2017) and Achilles (Tsherniak et al., 2017). In Achilles, the researchers used 501 cell lines and a genome scale library of 107,523 short hairpin RNAs (shRNAs). The Project DRIVE team opted for knocking down a smaller set of 7,837 genes across 398 cell lines, however they used an even larger 151,504 member library, with a median of 20 shRNA per gene. The authors of Achilles later published an improved, integrated version of their dataset by integrating DRIVE, Achilles, and Marcotte (Marcotte et al., 2016) data called DEMETER2 (McFarland et al., 2018).

In CaVu, we present the users the option to use either DRIVE alone or the DEMETER2 integration of DRIVE, Achilles and Marcotte. To assess the fitness of purpose of these datasets for our purposes, we used both to identify vulnerabilities for three separate diseases: carcinoma, melanoma, and hematopoietic & lymphoid neoplasm (Figure S1). We chose these diseases specifically so that they include both narrow (melanoma) and broad (H&L) disease definitions. We then used the Cancer Gene Census (CGC) to identify a list of established cancer drivers annotated as known drivers for each of these three diseases (Tate et al., 2019). We then assessed the number of CGC genes correctly identified as vulnerabilities for the appropriate cancer type. The results were better when using DRIVE for each of the three diseases. We used DRIVE when making the predictions validated in this study.

Here we emphasize that DEMETER2 has other significant advantages over DRIVE, the most important being its significantly enhanced coverage with both more genes and more cell lines. If the genes in the pathway hypothesized to be important for a particular disease are not among the 8k genes in DRIVE, then DEMETER2 is the only alternative. Likewise, if a disease has too few cell lines in DRIVE, which is the case for a number of diseases, then DEMETER2 is again the only choice available. Finally by incorporating DRIVE and Achilles, together with Marcotte (Marcotte et al., 2016), DEMETER2 leverages the strengths of each dataset. Therefore we would like to underline that both of these databases are very valuable, and hence we offer both DRIVE and DEMETER2 as options to the user.

### 2.3. Mapping vulnerabilities to drug candidates

There are two main motivations for identifying subtype specific vulnerabilities: understanding pathobiology and discovering therapeutic interventions. In order to expedite therapeutic discovery, we provide a list of chemicals targeting the identified vulnerabilities whenever available. We first use the Ensembl database to map the knockdown target genes to their protein products (Figure 1D). We then use the STITCH (Hubbard, 2002, Szklarczyk et al., 2015) database (v5) to map the proteins to the chemicals that interact with them. It is critical to ensure that the associations are high confidence. Therefore, we use only the protein chemical interactions that are supported with high confidence experimental evidence (high confidence means 70% or greater confidence, based on STITCH annotation). To enrich for chemicals more likely to be drug candidates, we restrict the molecular mass to be between 280 and 550 Daltons.

The users can potentially be interested exclusively in repurposing candidates. Approved drugs translate to clinic faster since they already have established safety profiles (Chong and Sullivan, 2007, Melnikova, 2012). To efficiently generate repurposing hypotheses, we annotate each chemical with similarity to an approved drug (Figure 1E). To do this, we first use the DrugBank dataset to retrieve the chemical structures of all drugs annotated as “approved” (2,502 unique chemicals) (Wishart et al., 2017). We then use RDKit (https://www.rdkit.org/) to calculate the similarity (Sørensen-Dice) of all the chemicals in our STITCH-derived list to all approved drugs. For each chemical, we then report the most similar known drug along with its chemical similarity. The user can quickly view this information and choose to prioritize the repurposing candidates or novel drug candidates. Alternatively, users interested in exploring novel chemistries can use this in the opposite manner, by searching for chemicals that are dissimilar to existing drugs.

### 2.4. Target identification validation

To test that our analysis can identify both known and novel vulnerabilities, we first searched for vulnerabilities of each tissue based lineage. We observed that our algorithm recovers well known lineage-specific driver genes (Figures 2A to 2F). For example, both SOX10 (Figure 2A) and MITF (Figure 2C) are transcription factors that are known to be drivers of melanoma, and both appeared as top lineage specific vulnerabilities for skin cancer. Beta-catenin (CTNNB1) is critical to APC-driven tumorigenesis and cell cycle regulation in colon cancer, and likewise it is identified to be the top colon cancer vulnerability (Figure 2B) (Morin, 1997, Tetsu and McCormick, 1999). Similarly, core-binding factor beta (CBFB) subunit, among other CBFs, is essential for haematopoiesis and plays crucial roles in leukemia, and CaVu accurately identifies CBFB knockdown to be selectively lethal for haematopoietic and lymphoid cancer cell lines (Figure 2D) (Speck and Gilliland, 2002). Using our reported statistical significance, users can filter out weaker hypotheses such GPR162 in kidney cancer (Figure 2G). Overall, there were 170,482 candidate lineage-specific vulnerability hypotheses, and our approach was able to accurately prioritize those hypotheses independently validated by the literature.

**Figure 2:**
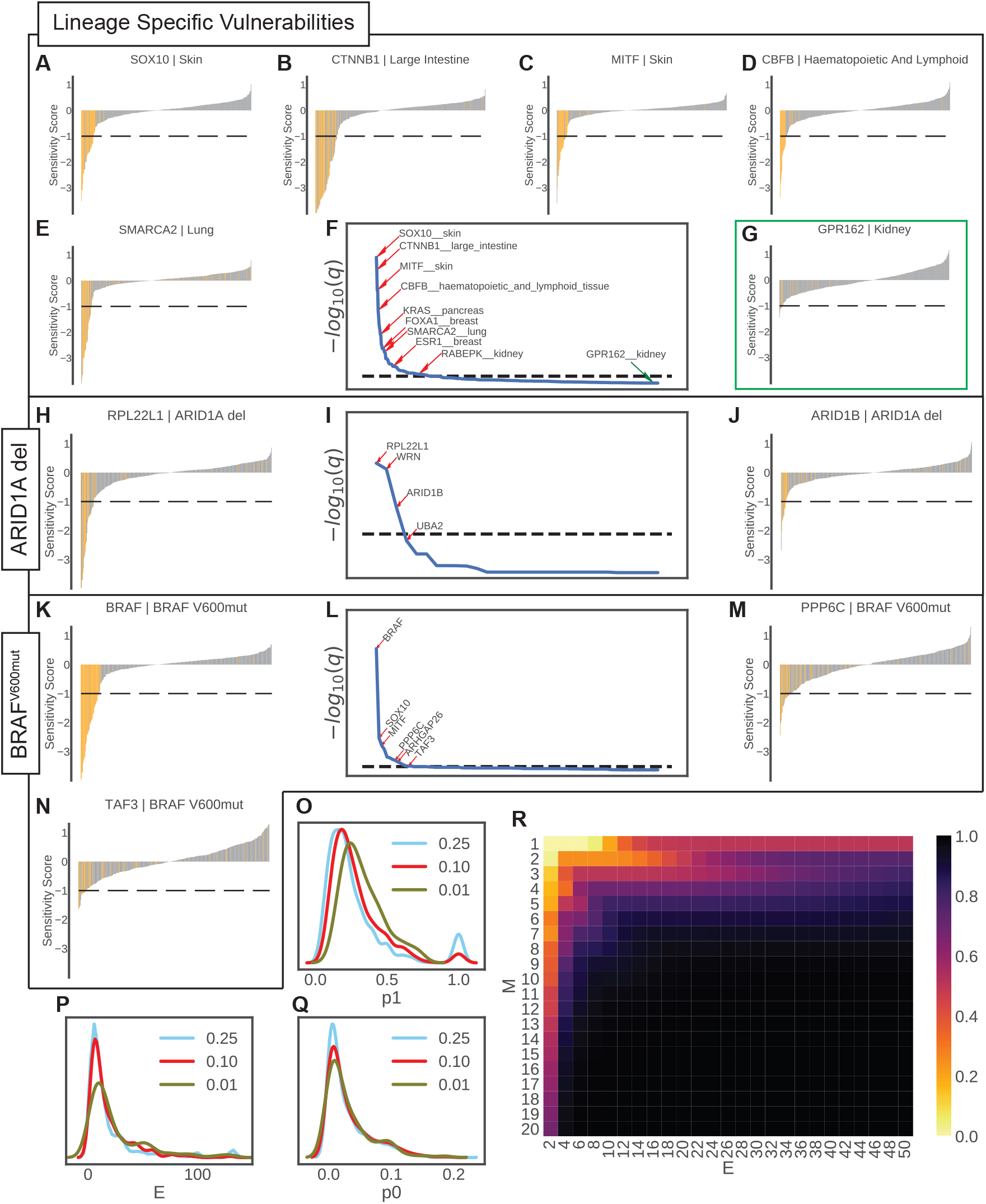
CaVu accurately identifies known vulnerabilities. To validate the vulnerability identification strategy, we evaluated the predicted vulnerabilities for both lineage-based (**A-G**) and mutation-based (ARID1A deleted, H-J; BRAF^V600mut^, **K-N**) analyses. In the sensitivity plots (**A-E, G, H, J, K, M, N**), the bars indicate the ATARiS sensitivity scores of knocking down a gene and the color of the bar indicates if the cell line is in the query set of cell lines (orange for query, gray for others). The top hypotheses for each query are reported (**F, I, L**) in sorted order, and the plotted hypotheses and some other significant hypotheses are indicated with red arrows. The knockdown of the statistically significant target candidates are visibly enriched for inducing lethality among cell lines in the targeted cancer subtype. One low-confidence hypothesis below the significance threshold is indicated with a green arrow and a green highlight (**G**), which expectedly lacks such enrichment. To identify the smallest number of cell lines for the subtype definition, we empirically established the likelihood of a real vulnerability inducing lethality among query cell lines (**O**), among non-query cell lines (**Q**), and effect size (**P**). We chose three different *q*-value cutoffs (0.01, 0.10, 0.25) to determine these empirical parameters at various levels of stringency, and observed that their means are rather stable. We used these parameters to run power analysis (**R**) and identified that having 7 cell lines in a user-defined subtype is sufficient for reasonable statistical power.

We further identified vulnerabilities for two molecularly defined subtypes of cancer: ARID1A deletion (31 cell lines), and BRAF^V600^ mutation (44 cell lines). Loss of ARID1A has been described to downregulate metastasis suppressors (Sun et al., 2017). Metastasis is an *in vivo* phenotype, whereas the data used in CaVu originates from 2D *in vitro* assays. Therefore, the ARID1A subtype definition tests whether CaVu can detect *in vivo* relevant phenotypic results. Indeed, CaVu detected significant vulnerabilities (Figure 2I), with the most significant being RPL22L1 (Figure 2H, *q*-value < 4.7e−6). However, even more surprising is that CaVu identifies in RPL22L1 a target that has been reported to promote metastasis by inducing epithelial-to-mesenchymal transition (EMT) (Wu et al., 2015). Although none of the experiments in DRIVE have been performed in the context of EMT, this finding suggests a mechanism whereby loss of ARID1A sensitizes cells to the loss of a known EMT driver. CaVu identified ARID1B as another significant (Figure 2J, *q*-value < 5.3e−4) vulnerability for ARID1A-deficient cancer, which was previously reported (Helming et al., 2014). CaVu also identified WRN as a significant vulnerability (*q*-value < 9.3e−6). However, the role of WRN in ARID1A-deficient cancer remains unknown, thus representing the potential for novel discovery. Overall, these findings illustrate the capacity of CaVu to prioritize hypotheses.

Among the vulnerabilities identified by CaVu for BRAF^V600^-mutant cancer (Figure 2L), the knockdown of BRAF itself was the most significant (Figure 2K, *q*-value < 6.9e−39). Given that BRAF^V600^ is a mutation leading to constitutive activation of the MAP-kinase pathway, this result served as a positive control (Ascierto et al., 2012). The next top two vulnerabilities were SOX10 (*q*-value < 1.6e−11) and MITF (*q*-value < 3.3e−9), which are known drivers in melanoma (Chong and Sullivan, 2007, Melnikova, 2012). BRAF mutations occur much more frequently in melanoma than other malignancies hence the enrichment of melanoma drivers (Davies et al., 2002). The next top vulnerability was PEA15 (Figure 2M, *q*-value < 5.7e−8) which blocks ERK-dependent transcription by anchoring ERK in the cytoplasm (Formstecher et al., 2001). CaVu enables us to discover that the function of this gene is essential for BRAF^V600^-mutant cancer suggesting a lethal relationship with ERK induced transcription for BRAF mutant cancer cell lines. Further down the list, DUSP4 (Figure 2N, *q*-value < 1.9e−5) is known to be promoted by BRAF activation of the MEK/ERK pathway, and associated with aggressive oncogenic behavior (Cagnol and Rivard, 2012, Wang et al., 2016). Other vulnerabilities with significant statistical evidence, such as ARHGAP26 (*q*-value < 3.7e−4) or TAF3 (*q*-value < 6.9e−3), have not been studied in the context of BRAF mutant cancer. Therefore, CaVu again both identifies known vulnerabilities that validate the model, and novel hypotheses that await future experimental exploration.

### 2.5. Power analysis

When composing queries, users would naturally need to know how many cell lines there should be in the set that fits their subtype definition in order to expect reasonable power. Clearly, a subtype with only one matching cell line is not likely to yield a meaningful analysis. To answer the question how many cell lines must be included in a subtype, we provide a power analysis.

The number of samples necessary to guarantee a specified power threshold depends on the effect size *E*. In general, the larger the effect size, the smaller the number of samples required. In this context, we define effect size as the ratio of two probabilities: the probability of a gene knockdown being lethal to a cell line within a vulnerable subtype (on-target lethality), defined as *p*_1_, and the probability that it is lethal to a cell line outside the subtype (off-target lethality), defined as *p*_0_. The effect size is then calculated as *E* = *p*_1_/*p*_0_.

To estimate the ranges for reasonable values, we calculated *p*_1_ (Figure 2O), *E* (Figure 2P), and *p*_0_ (Figure 2Q) empirically as ratios of lethal to non-lethal effects of knockdowns using lineage-specific vulnerability assessments above a *q*-value cutoff. We used three different cutoff values: 0.25, 0.1, and 0.01, which respectively yield the median *E* values of 9.95, 10.76, and 14.20. For the stringent cutoff of 0.01, the median value of *p*_1_ is 0.3 and therefore for the purposes of the power test, we set *p*_1_ to that value and calculated *p*_0_ using the specified value of *E*. We defined a range of reasonable values of *E* based on the observed values in our empirical evaluation. We decided that the reasonable range of the number of cell lines in the target set *M* is between 1 and 20. For values of *M* > 20, sufficient power is available to identify vulnerabilities with any moderate effect size.

For all reasonable *E* and *M* values, we calculated the power using simulation (Figure 2R; see Materials and Methods for details). We observe that when using 7 or more cell lines, the user can expect satisfactory power for almost all values of *E* that are commonly observed in our empirical evaluations. Therefore the power analysis suggests that users can define cancer subtypes containing as few as six or more cell lines and expect good predictive performance from CaVu.

### 2.6. Integrating the data into a unified model as a tensor

We initialized a tensor of drug-target-disease associations with an integrated confidence score for each component *T*_*ijk*_ derived from data. Specifically, the integrated confidence score for each component *T*_*ijk*_ = − log10(*q*_*jk*_)\**c*_*ij*_ where *q*_*jk*_ is the *q*-value of the hypothesized target *i*s significance for disease *k* based on data from Project DRIVE, and *c*_*ij*_ is the STITCH-reported confidence value for an interaction between the protein *i* and chemical *j*. The association score *T*_*ijk*_ represents the strength of the evidence that a particular drug *i*, through its interaction with target protein *j*, is effective against disease *k*. The remainder of the tensor is initialized as zero.

The tensor described herein can be used for two different types of predictions. If the diseases are defined as subtypes, then retrieving *d*^*^ = argmax_*i*_ *T*_*ijκ*_ would select the best drug *d*^*^ for disease κ. Furthermore, the tensor can flexibly be reorganized to admit different diseases (as described in Section 2.1) depending on user preferences. Alternatively, given a drug δ known to be effective in disease κ, the mechanism can be predicted by identifying the targets *t*^*^ = argmax_*j*_ *T*_δ*j*κ_. Therefore our representation is both flexible and useful for a variety of purposes.

### 2.7. Validation of CaVu predictions against melanoma

The best test of a drug discovery platform is to use it to find novel drug candidates. We therefore conducted an experimental validation of CaVu’s predictions. We used two separate definitions of disease: melanoma as a tissue lineage based disease definition, and ER+ breast cancer as a mutation based disease definition. We first discuss the melanoma drug candidate discovery process.

First, we selected eight approved drugs that were predicted to target at least one vulnerability specific to this tissue lineage, namely: bosutinib, crizotinib, dasatinib, ponatinib, sorafenib, pazopanib, nilotinib and sunitinib. All the repurposing and novel drug predictions were selected based on the integrated confidence score calculated as described in Section 2.6.

We selected these eight drugs such that they spanned the whole spectrum of scores (ranging from 509 for crizotinib to 13964 for sorafenib) with a bias towards those with high scores (Figure 3A). Although these drugs are used as antineoplastic agents in the clinic, none of them are indicated for any skin cancer. Thus, they constitute repurposing predictions. We then chose one novel drug candidate, SU11652, which has never been used or tested in the clinic for any indication, and was not previously reported to have activity against any skin cancer. For positive control, we used the current standard of care, clinical targeted therapy for BRAF^V600E^ mutant melanoma, which is a RAF/MEK inhibitor combination (RMIC) of 1 μM dabrafenib + 100 nM trametinib (Robert et al., 2015). Finally, as a negative control we included masitinib, which was predicted to target the haematopoietic and lymphoid lineage, but not skin cancer.

**Figure 3:**
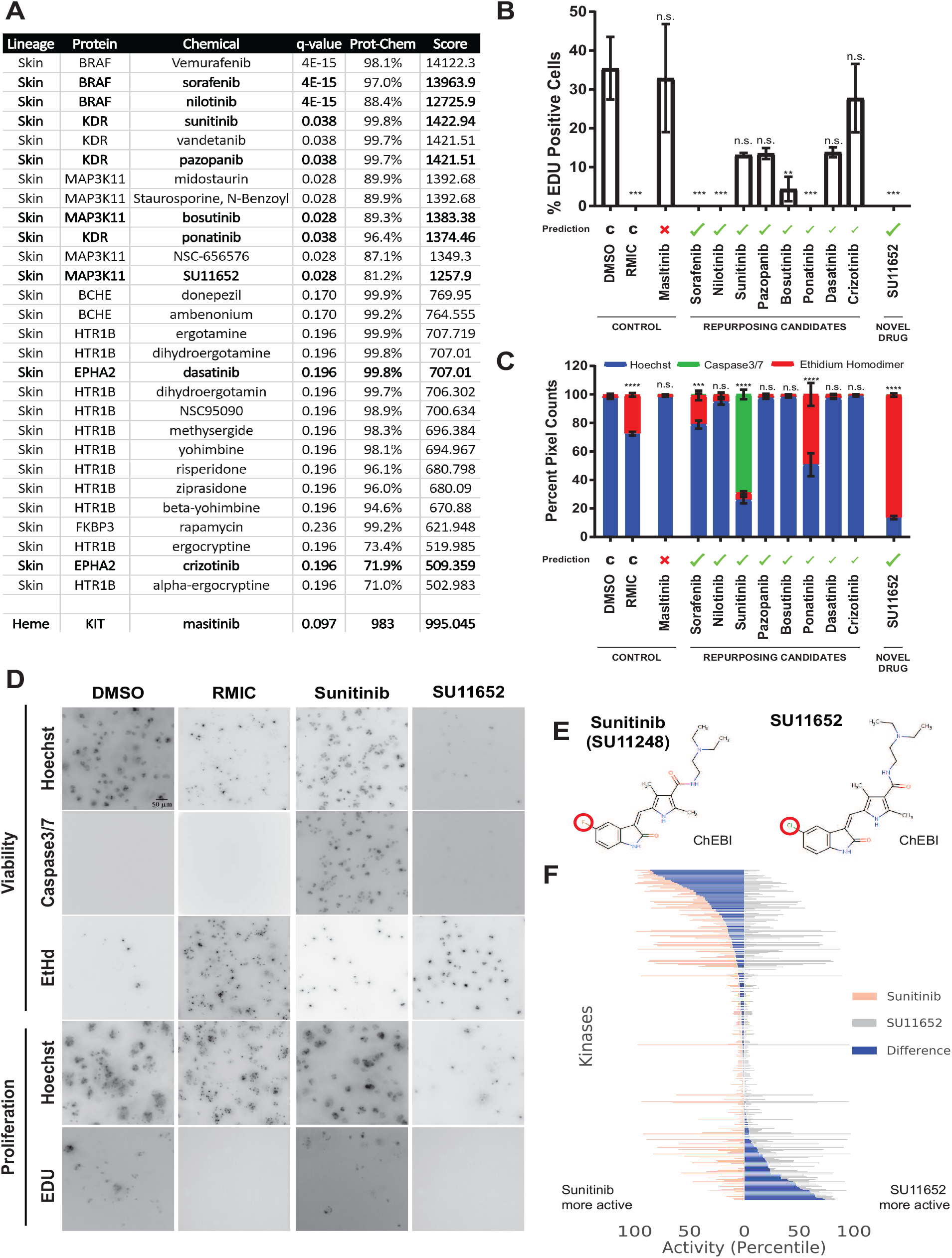
CaVu can successfully identify a novel melanoma drug candidate. Based on the confidence score calculated through CaVu, we chose 8 repurposing drugs and 1 novel drug candidate (highlighted in table) for skin cancer (**A**). A 10 μmol/dm^3^ screen to determine proliferation (**B**) and viability (**C**) was performed on A375 melanoma cells with these 9 drugs along with DMSO and Masitinib as negative controls and 1 μM Dabrafenib 100 nM Trametinib (RMIC) as a positive control. Results are shown as average ± SEM of three independent repeats with (\**p* < 0.05, \*\**p* < 0.005, \*\*\**p* < 0.0005). Representative maximum intensity projections of 3D image data (reduction of a volume onto a plane might look unfocused) showing viability and proliferation markers, with contrast measurements adjusted for visualization (**D**). Sunitinib and SU11652 are similar except for the highlighted fluorine / chlorine substitution (**E**). The difference between these two chemicals leads to significant bioactivity differences on 297 kinases based on data from Anastassiadis et al. with *p*-value < 1.4e−11 (Wilcoxon signed-rank test) (Anastassiadis et al., 2011). We show SU11652 (gray) and sunitinib (red) activity and the difference among them (blue) as percentile within the 178 kinase inhibitors tested, for each of the 297 kinases (**F**).

We tested the predicted drugs at a single dose, 10 μM, on A375 melanoma cells, which harbor the BRAF^V600E^ mutation. We chose this dose since the highest blood plasma concentration that can be achieved in the patients for a melanoma drug is 7.60 μM (Yamazaki et al., 2018), which means that if a chemical fails to achieve therapeutic response even at 10 μM then we can confidently rule out that compound as a therapeutic candidate. We used 3D collagen culture to create a physiologically relevant *in vitro* environment for drug testing, as described earlier (Murali et al., 2019). We used an imaging based readout of apoptosis, membrane permeabilization, and total viable nuclear area. We resolved overlaps among these different signals with a signal hierarchy: nuclear membrane permeabilization implies cell death and thus supersedes the other two, while apoptosis supersedes the viable nuclear signal. Then, for each treatment condition, we calculated the ratio of pixels from each channel to the total pixel count from all the channels. We determined cytostatic effects through imaging based *EdU* incorporation and Hoechst as a total nuclear marker. We quantified proliferation as the ratio of *EdU* to nuclear signal.

Four of the eight repurposing candidates (sorafenib, nilotinib, bosutinib and ponatinib), as well as SU11652 showed significant cytostatic activity (Figure 3B). Four out of these five compounds (sorafenib, ponatinib, sunitinib, and SU11652) also showed significant cytotoxic activity (Figure 3C). Note that the positive control RMIC had significant cytostatic and cytotoxic activity, whereas the negative prediction, masitinib, did not show any significant difference to the DMSO negative control, exactly as predicted.

Quite remarkably, SU11652 outperformed all other drugs, including the standard of care, RMIC. It caused 86% reduction in viability and stopped proliferation. RMIC has a strong cytostatic effect but a much weaker cytotoxic effect. Furthermore, SU11652 induced cytotoxicity through permeabilization within the 72 hour window. This is in contrast to the second most cytotoxic drug, sunitinib, which mostly had an apoptotic effect within the same time window (Figure 3D). This difference suggests that the cytotoxic effect of SU11652 progressed faster within the same time period. Furthermore, SU11652 completely prevented proliferation, which was not the case for sunitinib.

The difference in activity between sunitinib (i.e. SU11248) and SU11652 is caused by changes of only a few atoms, arguably the most significant being a fluorine to chlorine substitution (Figure 3E). We assessed whether if the similarity in structure implies similar bioactivity using the data from a 300 kinase activity screen (Anastassiadis et al., 2011). SU11652 and sunitinib are significantly different in their activity profile (Wilcoxon signed-rank test, *p*-value < 1.4e−11) over the 297 kinases for which both compounds have reported activity. The electronegativity difference between fluorine (in sunitinib) and chlorine (in SU11652) can potentially change the electron cloud across the larger molecule, and possibly explain this observation. Regardless of the cause, these two chemicals have large bioactivity differences despite seemingly few chemical differences as shown in Figure 3F using data from Anastassiadis et al. (2011). Therefore we decided that previous efforts that assessed sunitinib in the context of melanoma (Minor et al., 2012, Buchbinder et al., 2015) did not preclude the potential of SU11652 making a significant improvement. Due to the significant potential of SU11652, we decided to further characterize it as a melanoma drug candidate.

### 2.8. Characterization of SU11652 as a novel drug candidate against melanoma

We showed that SU11652 cytotoxicity generalizes to other models by testing it on four different melanoma cell lines (Figures 4A to 4D). We used two BRAF^V600E^ mutant (A375 in Figure 4A and M481 in Figure 4B) and two BRAF wild-type (BRAF^WT^) cell lines (M405 in Figure 4C and M498 in Figure 4D). We used RMIC as positive control for the BRAF^V600E^ mutants and trametinib alone for BRAF^WT^ melanoma cells. SU11652 had a dose-dependent cytotoxic effect on all four cell lines. We performed dose response assays for trametinib alone for the BRAF^WT^ cell lines and showed that it was largely ineffective (Figure S3). We also tested SU11652 on mouse melanoma models B16-BL6 (Figure S2A) and B16-F10 (Figure S2B) where SU11652 achieved an IC50 of 0.54 μM and 0.98 μM respectively. In conclusion, SU11652 was effective on multiple models of human and mouse melanoma, and more effective than the best available small molecule therapies.

**Figure 4:**
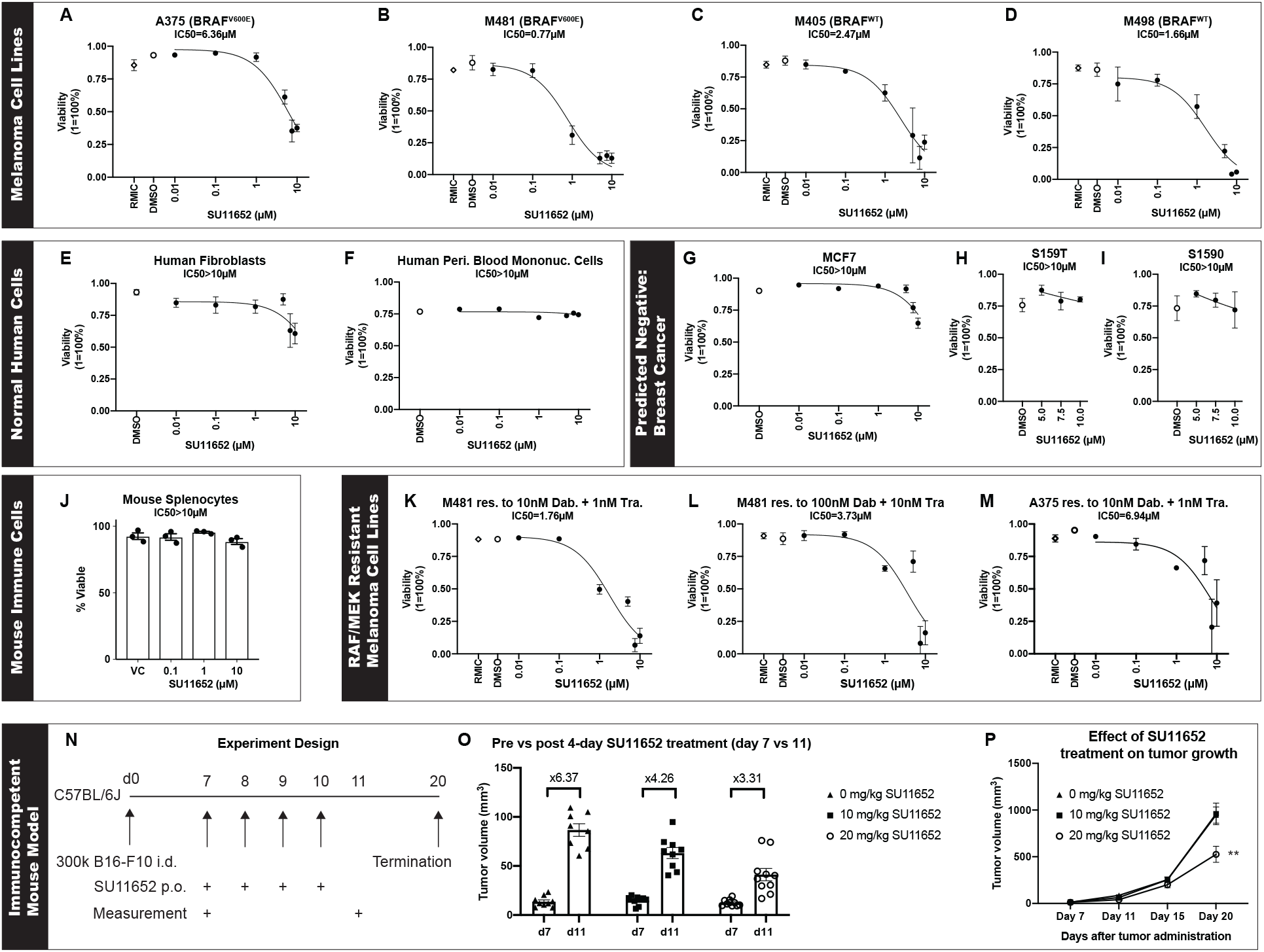
*In vitro* and *in vivo* validation of SU11652 as a melanoma drug candidate. *In vitro* dose response curves of two BRAF^V600E^ mutant cell lines, A375 (**A**) and M481 (**B**), and two BRAF^WT^ cell lines, M405 (**C**) and M498 (**D**) treated with SU11652. We used DMSO as negative control and RMIC (Raf-Mek Inhibitor Combo) as positive control for A375 and M481 cells. SU11652 was then validated on non-cancerous stromal cells: human foreskin fibroblasts 1 (HFF1) (**E**) and peripheral blood mononuclear cells (PBMCs) (**F**). All *in vitro* results are average ± SEM of three independent biological repeats. We validated a negative prediction, specifically that SU11652 would *not* have an effect on breast cancer using three different breast cancer cell lines: MCF7 (**G**), S159T (**H**), and S1590 (**I**). To demonstrate safety further, we showed that SU11652 does not cause any viability impairment on mouse splenocytes (**J**). We show that SU11652 can work on RMIC-resistant melanoma by applying SU11652 to BRAF^V600E^ melanoma which has been continously treated with RMIC until RMIC treatment ceases to impair growth (**K-M**). Finally, to demonstrate the *in vivo* efficacy of SU11652, we used a syngeneic mouse model of melanoma as an immunocompetent tumor model, and treated mice with SU11652 for only four days after the tumor was palpable at day 7 (**N**). We used increasing doses of SU11652 and measured tumor growth pre- and post-treatment (**O**). We then measured tumor volume until 20 days after tumor injection and showed that even a short 4-day treatment with SU11652 reduces tumor volume for more than two weeks after the end of treatment (**P**).

We then assessed whether the dose at which SU11652 shows activity is clinically plausible. In the clinic, the standard-of-care drug dabrafenib has maximum measured plasma concentration between the 95% confidence interval of 5.82 μM and 7.48 μM (Yamazaki et al., 2018). SU11652 shows *in vitro* activity at the 1 μM to 5 μM range, which compares favorably to the clinical dose observations.

To test the robustness of our findings, we also used a more established method of evaluating drug response, namely CellTiter-Glo (CTG) readout. Briefly, CTG reports relative chemiluminescence by measuring ATP activity, and is commonly used in drug discovery. SU11652 had similar effects on all four cell lines using the CTG assay (Figure S5). Therefore the impact of SU11652 was robust across different assay types.

Next, we tested for potential adverse toxic effects on non-malignant cells. Since the non-malignant cells in the tumor microenvironment are mostly stromal or immune cells (Tirosh et al., 2016), we used human foreskin fibroblasts (HFF1, Figure 4E), normal human immune cells (peripheral blood mononuclear cells; PBMCs, Figure 4F) and mouse splenocytes (Figure 4J). SU11652 did not have a significant cytotoxic effect on PBMCs or splenoctyes, and only a minor toxic effect on HFF1 at 10 μM. We tested for adverse effects on immune cell activation by treating splenocytes with SU11652 during anti-CD3/anti-CD28 stimulation. We chose this stimulation since it allowed us to specifically activate T cells, which is arguably the most important cell type for anti-tumor responses. SU11652 did not impact the viability of murine splenocytes after the 3-day culture period. Since SU11652 is active at the low micromolar range, this provides a good therapeutic window where the drug is cytotoxic against skin cancer cells but not normal cells from the human skin tissue.

Thus far, our experiments validated the absence of type I errors, namely whether our positive predictions were correct. Next, we tested for type II errors where CaVu missed activity that is indeed real. To that end, we identified breast cancer as one lineage where SU11652 was predicted to be inactive. We tested SU11652 on MCF7 breast cancer cell line (Figure 4G) as well as S1590 (Figure 4H) and S159T (Figure 4I) cell lines (Westcott et al., 2015). The IC50 of SU11652 was beyond the upper limit of detection range at 10 μM. This validated the CaVu prediction that SU11652 would not be active on breast cancer cell lines.

One major disadvantage of RMIC therapy on BRAF^V600E^ melanoma is the almost inevitable acquisition of resistance (Shaffer et al., 2017, Villanueva et al., 2011). Since CaVu predicted SU11652 on the basis of entirely different targets than BRAF or MEK, we hypothesized that SU11652 could be an effective treatment for BRAF and MEK inhibitor resistant (BRAFr/MEKr) melanoma cells. To generate BRAFr/MEKr melanoma cells, we continuously treated A375 or M481 cells with 10nM dabrafenib + 1nM trametinib or 100nM dabrafenib + 10nM trametinib for more than one month. We then conducted dose response viability assessments with SU11652 on the BRAFr/MEKr cells (Figures 4K to 4M). SU11652 was cytotoxic to the BRAFr/MEKr cells.

To test whether SU11652 activity transferred to the *in vivo* context of melanoma therapy, we designed the experiment shown in Figure 4N. Briefly, we injected 300k B16-F10 mouse melanoma cells into C57BL/6J mice. After the tumors were palpable at day 7, we injected SU11652 at 10 mg/kg or 20 mg/kg, or vehicle control for the next four days. We measured tumor volume until day 20. The tumor volume immediately before and after treatment showed a 6.3-fold increase in the vehicle control arm, which compares favorably to 4.26-fold for 10 mg/kg and 3.31-fold for 20 mg/kg (Figure 4O). We then followed the progression of the disease, despite giving no further drug injections, until day 20. There was a significant reduction in tumor volume even at day 20 in the 20 mg/kg dose arm (Figure 4P). Therefore SU11652 achieved durable control of tumor growth even with intermittent therapy.

The syngeneic mouse models have a normal immune system however their disease is mouse melanoma. To assess the impact of SU11652 on human models of melanoma, we performed another *in vivo* study. A375 (BRAF^V600E^) and M405 (BRAF^WT^ and NRAS^WT^) melanoma cells were injected in NOD-scid IL2rg^null^ (NSG) mice. Once palpable the mice were treated with vehicle control, sunitinib and SU11652 at the 20 mg/kg dose. For the A375 model, we included an additional arm of dabrafenib and trametinib combination therapy as a positive control. For the BRAF^WT^ M405 model, we omitted such an arm simply because there are no established effective treatments for BRAF^WT^ melanoma. As we showed in Figure S3, trametinib alone is not effective. In a relatively recent phase II trial that assessed trametinib in combination with a novel Akt inhibitor (GSK2141795) in BRAF^WT^ and NRAS^WT^ melanoma (Algazi et al., 2017) there was no significant clinical activity. Due to the absence of a rationale for a treatment arm, we chose to omit sacrificing mice for a “positive control” arm.

SU11652 controlled tumor growth in comparison to both vehicle control and sunitinib in both models (Figure S4). Furthermore, the mice treated with SU11652 survived longer than sunitinib treated mice before they had to be euthanized. The weight of the SU11652 treated mice were similar to the vehicle control indicating no gross adverse effect on mice as shown in Figures S4C and S4D. In the A375 melanoma model, the dabrafenib and trametinib RAF/MEK inhibition combo (RMIC) therapy was clearly more effective than SU11652, which is to be expected since this is a BRAF^V600E^ melanoma model. However, RMIC treatment eventually leads to resistance and, when that happens, SU11652 still remains effective as we show in Figures 4K to 4M. Therefore we determine that SU11652 is a potential candidate for first-line treatment of BRAF^WT^ melanoma and second-line treatment of BRAF^V600E^ melanoma.

### 2.9. Low rank tensor completion based mechanism prediction

To predict the mechanism of action of SU11652, we developed a machine learning pipeline that can predict the most likely novel drug-target-disease associations. We represent the tripartite drug-target-disease network as a third order tensor with drugs, targets, and diseases used to index association scores (calculated as described in Section 2.6). The drug-target-disease tensor is extremely sparse: defining diseases based on the tissue of origin, only 0.02% of the entries are known. Completing the remaining 99.98% from such a small number of known entries is a challenging problem, however, it has been well studied for over a decade because most technology companies have to complete similarly large and sparse tensors (ex: user-movie-time tensor for Netflix) to make recommendations to their users (Xiong et al., 2010).

The common strategy when completing such large tensors on the basis of sparse known entires is to assume the tensor is low rank and complete it on the basis of this assumption, a field known as low rank tensor completion (LRTC). Likewise, here we also assume the tensor is low rank, perform CANDECOMP/PARAFAC (CP) tensor decomposition to find a lower dimensional latent representation that encodes the associations, and reconstruct the tensor from that representation. CP decomposition represents an input tensor as a sum of *R* rank one outer products, with *R* being the rank of the decomposition. CP decomposition has early theoretical roots in mathematics (Hitchcock, 1927), relatively early computational implementations (Harshman, 1970), and remains a foundational LRTC method that is still actively used (Long et al., 2019). However to the best of our knowledge, this is the first time it is applied to drive cancer drug discovery predictions.

To assess the predictive power of LRTC in the context of a cancer-specific drug-target-disease tensor, first we ran a simulation by sequestering known data (Figure 5A). We randomly withheld 0.05% of eligible tensor components, performed LRTC, and chose the highest scoring interactions as the predictions. We then set a budget of 100 predictions and created two modes of LRTC: an “active learning” variant that we retrained after testing each batch of five predictions, and a “passive learning” variant that used the same list of predictions for the entire budget. In this, we followed a classical definition of active & passive previously used in literature (Warmuth et al., 2003). We considered a prediction a “hit” if and only if it was a known entry that was held out. This is strictly an underestimate because there can be novel predictions that would be hits if tested, which is the entire point of LRTC, yet these are counted as misses in this *in silico* validation. Nevertheless, this is still useful for assessing overall applicability of LRTC in this problem domain. We estimated the hit rate that would be observed with brute-force high throughput screening via the expectation of random guessing. We then calculated the “learning factor” as the improvement over random as the ratio of the hits found via LRTC to random hits. We repeated this entire procedure 16 times to remove the effect of randomness in the data hold-out phase. Our model, in both the active and passive variants, consistently demonstrated > 20% hit rate over 100 predictions and 3,000 to 4,000 fold improvement over random, showing that LRTC has strong predictive potential in a cancer tensor.

**Figure 5:**
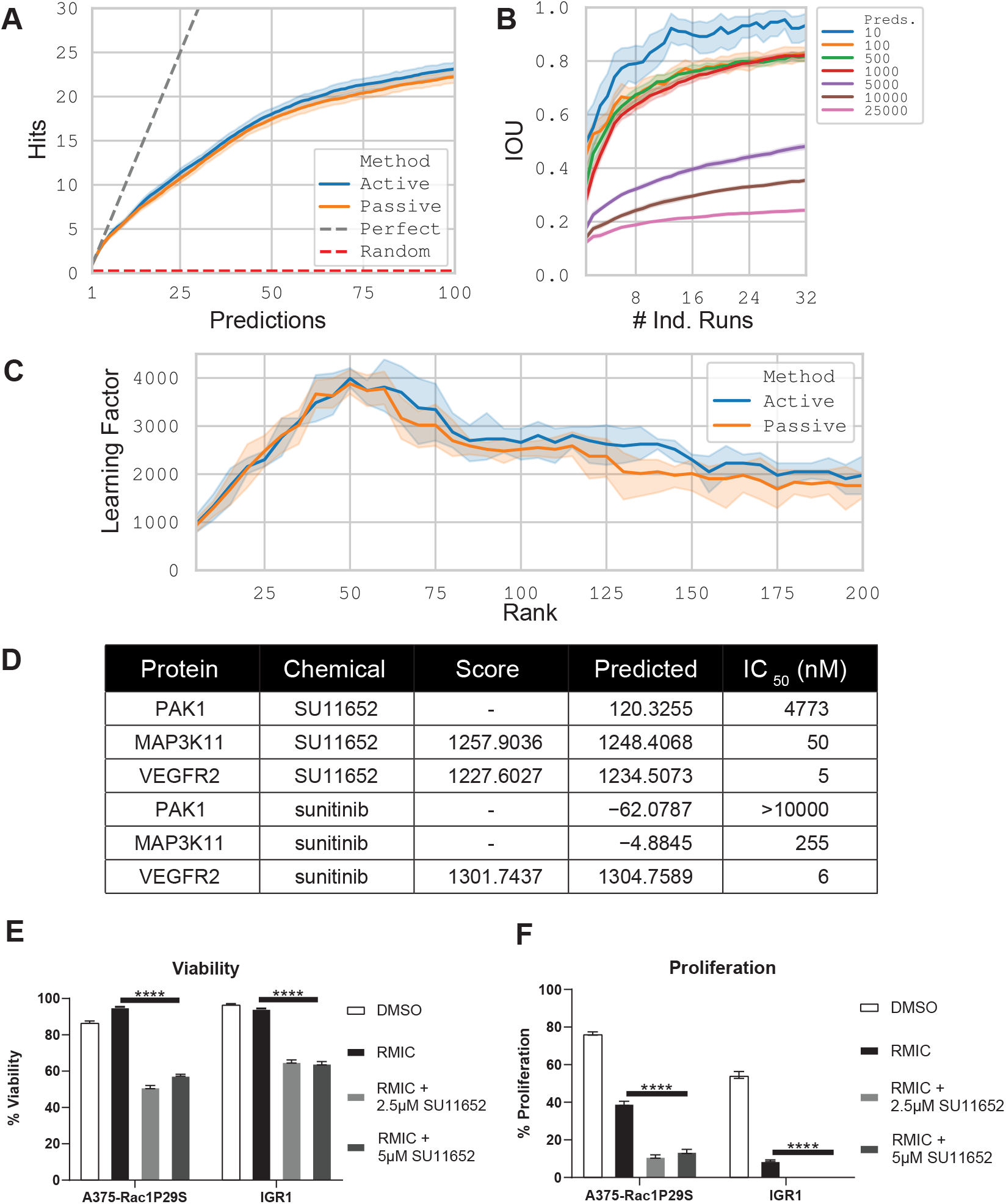
LRTC accurately predicts mechanism of action of SU11652. To predict the mechanism of SU11652, we constructed a third-order tensor of drug-target-disease associations and showed that LRTC can recover known entries removed from the tensor (**A**). To assess model robustness, we compared the agreement (intersection over union; IOU) among the top predictions (ranging from 10 to 25,000) for two separate runs with the same number of models in each. Increasing numbers of models increase the robustness, yet there is diminishing returns after 16 (**B**). The drug-target-disease tensor is likely low rank because using more or less than 50 ranks in the decomposition degrades model performance (**C**). We used LRTC to predict the unknown target of PAK1 for SU11652, in addition to two other targets reported in STITCH (MAP3K11 and KDR), and tested their activity *in vitro* to measure the IC50 of inhibition by SU11652 (**D**). Sunitinib was predicted *not* to interact with PAK1 and likewise found not to target PAK1. The novel associations did not have any scores before the tensor decomposition, hence their entry in the score column is simply a dash. The RAC1^P29S^ hyperactivating mutation drives proliferation through multiple pathways only one of which is PAK1 dependent and therefore confers resistance to SU11652 in RAC1^P29S^ knock-in A375 and the RAC1^P29S^ cell line IGR1 as measured by both viability (**E**) and proliferation (**F**).

The initialization of the CP decomposition can, as in most machine learning models, affect the trained model and thereby the performance. To counteract this, we initialize a multitude of independent models and use the consensus predictions. We performed a comprehensive analysis of prediction robustness (as measured by intersection over union; IOU) when using different numbers of independent initializations. The number of predictions to be considered is an influencing factor in this assessment, and therefore we used a range of values from 10 to 25,000 predictions. In practice, we expect only the lower end of this range to be relevant for two reasons. First, it is hardly feasible to test 25,000 drug-target-disease hypotheses. Second, we expect an active learning strategy in practice where the model is retrained after testing a batch of predictions. We observed that our predictions are generally stable when using more than 10 initializations, and consistently so when using 16 or more (Figure 5B). Among the top scored components between two independent 16-initialization models, at least 9 are shared out of 10 predictions with high confidence. Therefore the predictions are generally robust when models are properly initialized.

We then assessed the applicability of the low rank assumption for the tensor. CP rank of a tensor is defined as the rank *R* at which CP decomposition yields a perfect reconstruction. Computing the actual rank of the tensor requires observing all the entries in the tensor; hence it is impossible to know the real rank of our cancer-specific tensor with only 0.02% observed entries. However we hypothesized that using *R* lower or higher than the actual CP rank of the tensor should degrade completion performance. This is because lower than real rank should lead to models with insufficient descriptive capacity and higher than real rank should lead to overfitting. We observed a peak of performance around rank 50 (Figure 5C), suggesting that the actual rank of the tensor is likely to be low.

Having established a novel machine learning method for predicting drug-target-disease associations, we predicted the most likely yet unknown target of SU11652 against the “melanoma” disease as PAK1 (Figure 5D). Alternatively, this can also be viewed as identifying the index of the highest scoring target in the SU11652-melanoma fiber of the tensor. Interestingly, PAK1 was not predicted to be a target of sunitinib. We also observed that two other kinases, namely VEGFR2 and MAP3K11, were already present in the tensor with high scores. We used the EuroFins KinaseProfiler™ and IC_50_Profiler™ services to measure the activity of both sunitinib (SU11248) and SU11652 against these three kinases (Figure S6). SU11652 was found to inhibit all VEGFR2 and MAP3K11 with an IC_50_ in the nM range and PAK1 at IC_50_ = 4.7 μM; sunitinib was, as predicted, found not to have an effect on PAK1.

PAK1 is a complicated melanoma target. PAK1 is “melanogenic” in normal melanocytes and oncogenic in melanomas (Be Tu et al., 2017). It is recognized as a target in both BRAF^WT^ (Ong et al., 2013) and RMIC-resistant BRAF^mut^ melanoma (Babagana et al., 2017). However, the Pfizer drug PF-3758309 which targets all PAKs with nanomolar potency, including PAK1 at K_i_= 13.7 ± 1.8nM, (Murray et al., 2010) failed in a clinical trial on patients with advanced solid tumors at Phase 1 (NCT00932126). The early termination was attributed to undesirable PK characteristics and lack of an observed dose-response relationship. PAK1 is involved in a complex network of interactions with roles as diverse as cytoskeletal remodeling and PI3K activation, with a multitude of physiological roles (Kichina et al., 2010). As a result, a relatively weak inhibition of PAK1 by SU11652, at 4.7 μM, can contribute to its potential as a drug candidate by having enough activity to inhibit PAK1 activity in the cancer cells while not being strong enough to induce toxicity.

PAK1 and MAP3K11, both targets of SU11652, have the Cdc42/Rac interactive binding (CRIB) domain and therefore they are downstream effectors of RAC1 (Krauthammer et al., 2012). The hyperactive RAC1 P29S variant, the third most common mutation in melanomas, stimulates proliferation and mesenchymal-like transformation through the AKT, MRTF/SRF, and PAK pathways (Lionarons et al., 2019). Furthermore, another recent study showed that after RMIC treatment, RAC1^P29S^ can induce proliferation through the phosphoinactivation of NF2 in subcellular microdomains, also mediated through PAK1 (Mohan et al., 2019). Based on this multitude of compensatory pathways, we hypothesized that the PAK1 and MAP3K11 inhibiting SU11652 should at best be partially effective at controlling proliferation of RAC1^P29S^ melanoma after RMIC treatment. We tested this hypothesis using RAC1^P29S^ knock-in A375 cells (which are normally RAC1^WT^ and controlled by SU11652) and IGR1 which is a RAC1^P29S^ melanoma cell line. SU11652 was effective at reducing both viability (Figure 5E) and proliferation (Figure 5F) compared to RMIC, however with a diminished effect compared to its activity on RAC1^WT^ models (Figures 4A to 4D).

### 2.10. CaVu identifies the ER-positive breast cancer drug candidate TC-E 5008

To demonstrate that our methodology can be readily applied to an entirely different cancer subtype, we used CaVu to predict drug candidates against ER-positive breast cancer. We selected three drug-target-disease triplets for experimental validation on three cell line models: “BMS309403— FABP5” (Figure S7A), “Z1445513748—KDM1A” (Figure S7B) and “TCE5008—IDH1” (Figure 6A). Here, we were able to search over both drug *and* target hypotheses jointly due to our tensor based computational hypothesis selection engine. TC-E 5008 clearly outperformed the other two drug candidates, therefore we selected it for further validation. To test this molecule’s specificity against the ER-positive subtype of breast cancer, we also tested it against a triple negative breast cancer cell line where it demonstrated little to no activity (Figure S7C). TC-E 5008’s activity is likely due to a polypharmacological effect and not just IDH1, similar to SU11652, since two other IDH1 inhibitors did not have the same activity (Figure 6B).

**Figure 6:**
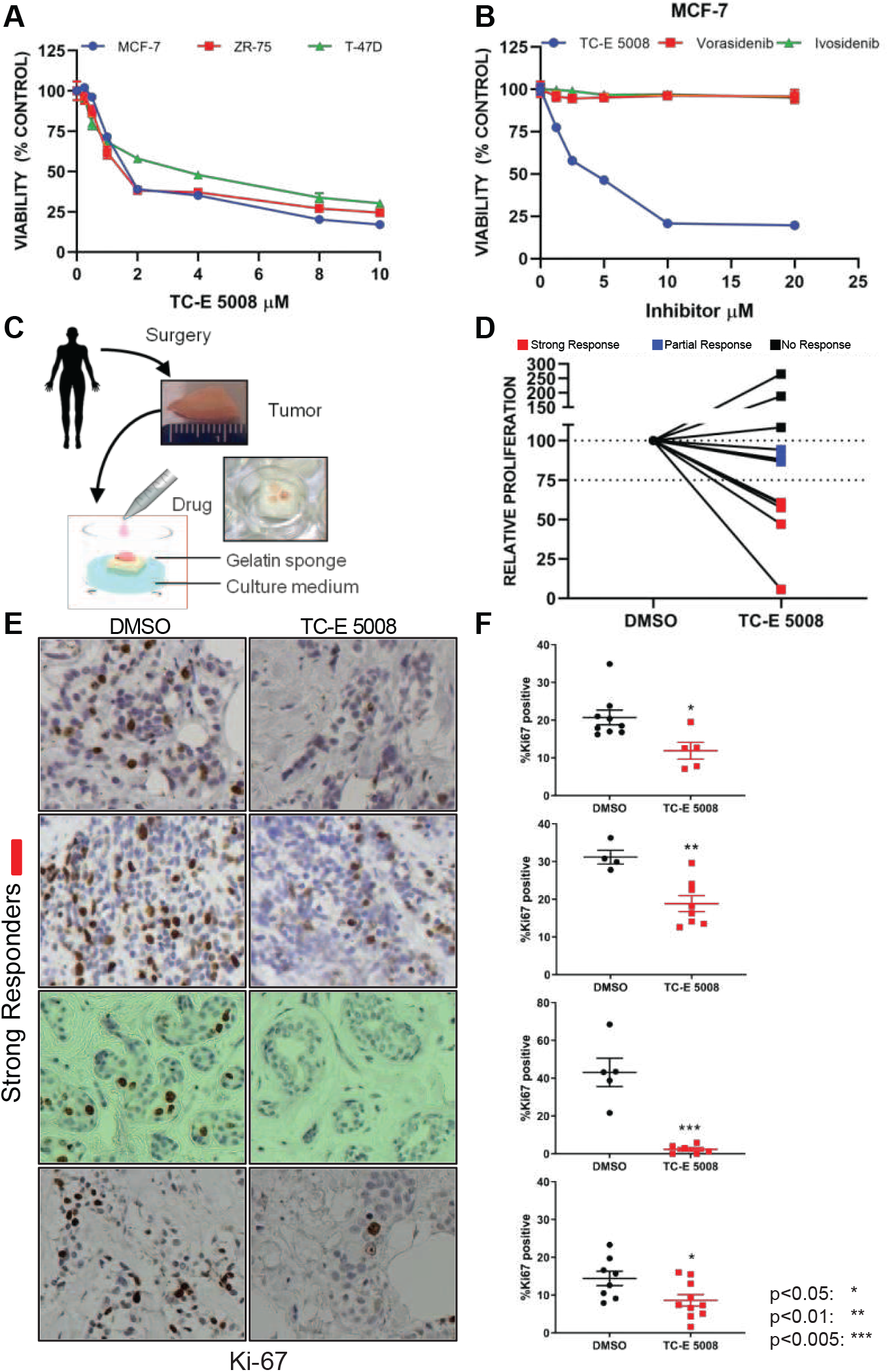
*In vitro* and *in vivo* validation of TC-E 5008 as an ER+ breast cancer drug candidate. TC-E 5008 has anti-proliferative activity on multiple ER positive breast cancer cell lines as measured by the CellTiterGlo assay (**A**). TC-E 5008 has differential activity compared to other mutant IDH1 inhibitors vorasidenib and ivosidenib on MCF-7 cells (**B**). Schematic representation of patient-derived explant (PDE) model with 72h drug treatment (**C**). TC-E 5008 treatment (10 μM) led to strong response (< 75% reduction in proliferation) in tumor samples from 4 patients, partial response (> 75% and < 100% proliferation) in 3 patient samples and no reduction in 3 patients (**D**). One representative image from each strong responder (**E**). Proliferation signal in all independent fields of view for each strong responder, with indicated mean ± SEM, and statistical significance relative to DMSO (**F**).

We validated the activity of TC-E 5008 on ten patient samples using our *ex vivo* human tumor culture assay, which allows for the validation of drugs on breast tumors in their native tissue architecture (Schiewer et al., 2012, Shafi et al., 2018). In brief, surgically resected breast tissues are sliced into small pieces and grown *ex vivo* for short term on a gelatin sponge in the absence or presence of desired compound (Figure 6C). In order to test TC-E 5008 in a variety of ER-positive tumors, we only used estrogen receptor positivity as inclusion criteria for the breast tumor samples. Incubation of TC-E 5008 with ER-positive breast tumor samples decreased their proliferation (Ki-67 staining) in 7/10 patients with 4/10 showing a significant reduction ranging from 40-95% (strong responders) and 3/10 showing a reduction in Ki-67 staining between 0-25% (partial responders) as shown in Figures 6E and 6F. Three tumor tissues did not show a decrease in proliferation (non-responders) when compared to untreated controls (Figure S8). We did not observe major morphological changes in the patient tissues treated with TC-E 5008 when compared to controls.

We anticipated a range of anti-proliferative activity of TC-E 5008 since patient stratification was kept to a minimum. In this limited cohort a 40% response rate suggests that TC-E 5008 has the potential to impact the growth of human breast tumors expressing ER as predicted by CaVu. A larger cohort will be necessary to identify which patient population is more likely to respond to TC-E 5008 and the overall efficacy in ER-positive breast tumors.

## 3. Discussion

With this study, our main contribution is to describe and validate a novel approach to cancer drug discovery. In our algorithmic therapeutic oncology paradigm, joint computational analysis of heterogeneous public datasets creates a tripartite drug-target-disease interaction network. This network is then converted into a third order tensor and completed using low rank tensor completion. The resultant tensor enables rapid and effective search of new therapeutic candidates by jointly searching over the best drug and target combinations for each separate cancer (sub)type.

We extensively validate our methodology by testing predictions against two separate diseases, melanoma and ER-positive breast cancer. We use a multitude of assays and readouts ranging from CellTiterGlo to fluorescent multi-readout high content assays to *ex vivo* human tumor based assays. We show that we are able to find effective drug candidates, as well as develop meaningful hypotheses about the mechanism of at least one of these drug candidates, SU11652. We computationally validate our approach through LRTC simulations with held-out data, perform literature validation of predictions, and present a power analysis to serve as a guide to define diseases. Finally, we provide a web server (accessible at https://cavu.biohpc.swmed.edu/) that serves to democratize our methodology to the entire community of cancer researchers.

As with any study, there are certain limitations. Any limitations in STITCH or DRIVE will clearly impact the results of CaVu. However, these datasets are continually updated. Curators constantly improve the quality of drug-target interactions reported in STITCH, which is in its fifth version. Similarly governmental and private research institutions are likely to conduct and publish target identification work such as DRIVE. Continued work in this area is probably going to transition from knockdown to knockout technologies using CRISPR. The computational framework we present here is highly modular, and therefore it can accommodate new versions of these data sources seamlessly. We designed the web server, CaVu, to likewise be highly modular and plan to regularly update it as major revisions of the underlying data sources are released. Therefore, we envision CaVu as an ongoing discovery portal that improves along with its data sources.

Our results with SU11652 demonstrate that already in the present state of CaVu there are good reasons to use this tool as an automated hypothesis discovery engine. It enables rapid acquisition of novel, unbiased, data-guided hypotheses. These hypotheses have significant translational potential. By integrating the normally disparate stages of drug discovery—namely target identification, hit identification, and lead selection—we enable efficient progress to translation. The efficiency with which CaVu can yield translational hypotheses with small numbers of chemicals can enable the use of clinically relevant but complicated assays such as *ex vivo* tumor culture, as we demonstrate with TC-E 5008. *Ex vivo* human tumor based assays, with a reported 87% clinical translation accuracy (Majumder et al., 2015), would produce therapeutic candidates that are more likely to impact clinical practice.

In the future, our work can be extended in multiple directions. Currently, we only rely on known drug-target interactions as reported by STITCH. However, it is possible to develop an improved version of CaVu that incorporates known and predicted drug-target interactions. This improvement would render a larger percentage of the identified vulnerabilities actionable. Furthermore, when a drug candidate looks promising, such as SU11652, there are often follow-up questions regarding its mechanism and target. SU11652 outperformed standard-of-care therapy for two of the five models, showing heterogeneity in response as observed by most cancer therapies. These results prompt further investigation to identify the molecular characteristics of the responders as well as non-responders. Furthermore, computational predictions to answer mechanistic queries can potentially be built into CaVu. Similarly, it is possible to incorporate computational work for vulnerability identification based on gene expression and thus extend to other cancer subtypes for which expression data exists but knockdown data does not. More broadly, we hope that our work becomes a platform for bridging the bioinformatics and cancer research communities, and help make cancer drug discovery more efficient.

## 4. Materials and Methods

### 4.1. Vulnerability assessment for a molecularly defined cancer subtype

We used the hypergeometric enrichment test to identify gene knockdowns that cause enriched lethality for specific lineages. Given that lineage *l* has *c*_*l*_ unique cell lines, knockdown of gene *g* is observed to be lethal for *t*_*g*_ cell lines, and the total number of unique cell lines is *T*, let 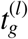 be the number of cell lines within lineage *l* for which the knockdown of *g* is lethal. We assume that 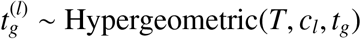 where:

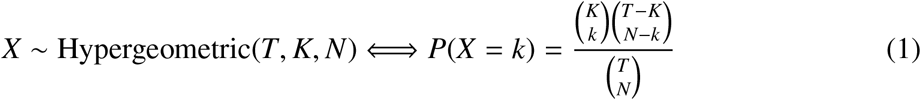

To calculate the enrichment *p*-value, we calculated 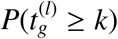 where *k* is the observed number of lineage *l*-specific cell lines for which knockdown of gene *g* is lethal. Since we used the ATARiS quantification method (Shao et al., 2012) based analysis of DRIVE which yielded 6,557 genes with reliable knockdown on 398 cell lines, we calculated 2.6 million such *p*-values, which strongly requires multiple hypothesis correction. We applied Benjamini-Hochberg false discovery rate (FDR) correction (Benjamini and Hochberg, 1995). We found 23,387 gene-lineage pairs that met an FDR threshold of 0.25, while only 626 pairs were below the 0.1 FDR threshold.

### 4.2. Linking vulnerabilities to drug candidates

We used the STITCH 5 database (Szklarczyk et al., 2015) to link proteins to the small molecules known to target them. We used only the experimental interaction confidence channel because it is the most reliable source of information. Furthermore, we employed a stringent confidence cutoff of 70%, thereby leaving only the interactions that STITCH annotates as high confidence or above. These molecules contained many chemicals that are inadmissible as drug candidates, and thus required filtering for drug-likeness.

Drugs that are highly similar to existing drugs are, by definition, drug-like. Therefore we used the DrugBank v5 (Wishart et al., 2017) to retrieve the list of small molecule drugs annotated as approved for pharmaceutical use. We then used RDKit greater similarity to any approved drug. Finally, 7,934 unique chemicals were found to target a lineage-specific vulnerability gene (with FDR cutoff of 0.25) with high confidence (70% or greater STITCH confidence) and also have a drug-like structure.

### 4.3. Cell Lines

A375 melanoma cells harboring the BRAF^V600E^ mutation were purchased from ATCC (CRL-1619). Patient derived xenograft (PDX) cell models M405, M481 and M498 were acquired from the University of Michigan via Sean Morrison at UT Southwestern Medical Center. M481 harbors a BRAF^V600E^ mutation. M405 and M498 are BRAF wild type (WT) melanoma cell lines. Human foreskin fibroblast 1 (HFF1) was purchased from ATCC (SCRC-1041). MCF7 breast cancer cells were purchased from ATCC (HTB-22). Human breast cancer cells ZR-75, T-47D, MDA-MB-231 were either obtained from American Type Culture Collection (ATCC, Manassas, VA) or a kind gift from Dr. John Minna at UT Southwestern. S1590 and S159T breast cancer cells were acquired via Gray W. Pearson at UT Southwestern Medical Center. Each aliquot of the cells is used no more than 20 passages after which a fresh aliquot is used. Mycoplasma testing was performed using Mycoscope, a PCR based mycoplasma detection kit. Mycoplasma testing was done for almost every 6 to 9 months. No mycoplasma detected in any of the cell lines used in the manuscript. Human peripheral blood mononuclear cells were isolated from three de-identified buffy coats acquired from Carter Blood Care.

### 4.4. Determination of tensor rank

To support our hypothesis that *de novo* drug discovery can be formulated as a low-rank tensor completion problem, we performed multi-sampled CP decomposition with varying ranks of the latent space using the TensorLy software package (Kossaifi et al., 2019). Computations were distributed across the UT Southwestern BioHPC using Ray (Liaw et al., 2018).

We used 16 different train-test tensor splits for each tested rank. Components moved from training to testing were restricted such that their removal would not result in a zero vector for any previously nonzero vector across any dimension. Thus, no elements of the test set would require the model to guess with no information. Each of the 16 individual tensor completions were initialized with different random seeds, but the seeds were kept constant across all trials to ensure comparable results. Both active and passive performance peaked at rank 50 before trailing off.

### 4.5. Cell line count power analysis

To estimate the number of cell lines to have in each subtype description to have statistical power, we performed a simulation method. First, we recognize that even genes that are vulnerabilities for a particular subtype might not necessarily be lethal for all cell lines in that group. The probability that a gene whose knockdown is lethal for a subtype being lethal for any single cell line in that subtype is defined as *p*_1_. Conversely, we also recognize that the genes that are not subtype-specific vulnerabilities for a cell line can nevertheless be lethal. We define *p*_0_ to be the probability of a gene knockdown being lethal for any single cell line defining a target subtype even though the gene is not a subtype specific vulnerability. We then define the effect size (*E*) in terms of these two probabilities defined as follows: *E* = *p*_1_/*p*_0_. Empirically, we found *p*_1_ = 0.3 to be a reasonable estimate (Figure 2O). Therefore for any given *E* value, we calculate *p*_0_ = *E*/*p*_1_. We define the number of cell lines that comprise the target subtype as *M*. We set the total number of cell lines not in the target set trivially to *N* − *M* where *N* is the total number of cell lines. We conducted the simulations over a range of *E* values from 2 to 50 and *M* values from 1 to 20. For a given set of values for *E*, *M*, *N*, we sampled 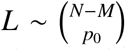 which is the total number of cell lines outside of the query set (i.e. those cell lines that are not in the target subtype) for which a gene knockdown is lethal. We also sampled 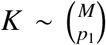 as the total number of cell lines that are in the query set (i.e. those cell lines matching the target subtype definition) for which a particular gene is lethal given that the gene is a subtype-specific vulnerability. We then calculated the hypergeometric enrichment test we use for our target identification, with a *p*-value cutoff of 0.05 to calculate power. We repeat this process 100 million times, and report the results in (Figure 2R).

### 4.6. Cell Culture Materials

A375 cells were cultured in Dulbeccos modified eagle media (DMEM), with high glucose and L-glutamine (11965-167) was purchased from Thermo Fisher Scientific, supplemented with 10% fetal bovine serum (FBS), also from Thermo Fisher Scientific (11965-167). M405, M481 and M498 melanoma cells were cultured in dermal basal medium purchased from ATCC (PCS-200-030), supplemented with adult melanocyte growth kit purchased from ATCC (PCS-200-042), which includes rh insulin, ascorbic acid, L-Glutamine, epinephrine, CaCl2, peptide growth factor and M8 supplement. S1590 and S159T cells were cultured in F-12 base medium supplemented with 5% FBS, 5 μg/mL bovine insulin (Sigma I1882) and 1 μg/mL hydrocortisone (Sigma H0888). Phenol red free DMEM with high glucose and L-glutamine (21063-045) for fluorescence imaging was purchased from Thermo Fisher Scientific. The phenol red free media was supplemented with 10% FBS. Trypsin/EDTA (R001100), trypsin Neutralizer (TN) (R-002-100), Dimethyl Sulfoxide (DMSO) (BP231-100) were purchased from Thermo Fisher Scientific. For 3D culture, bovine collagen I with a molecular weight of 300 kDa was purchased from Advanced Biomatrix (5005-100). Dabrafenib (GSK2118436), and Trametinib (GSK1120212), Pazopanib (S3012), Nilotinib (S1033), Sorafenib (S7397), Dasatinib (S1021), Ponatinib (S1490), Vorasidenib (S8611), Ivosidenib (S8206) and Masitinib (S1064) were purchased from Selleckchem. Bosutinib (B1788) and sunitinib (S8803) were purchased from LC laboratories. SU11652 (sc-204310) was purchased from Santa Cruz Biotechnology. TC-E 5008 (5244) was purchased from Tocris Bioscience. BMS-309403 (BM0015) was purchased from Sigma. Z1445513748 was purchased from Enamine. We bought the following from Thermo Fisher Scientific: the cell impermeant viability marker ethidium homodimer (E1169); CellEvent, a caspase 3/7 (C10423) green detection reagent to identify apoptotic cells; Hoechst 33342, nuclear stain (H3570); and Click-iT based imaging kit (C10340). CellTiter-Glo was purchased from Promega for 3D (G968A) assays. Human interleukin-2 (IL-2) (H7041) was purchased from Sigma Aldrich. Ficoll-PaqueTM PLUS (17-1440-02) to isolate PBMCs from buffy coats was purchased from GE Healthcare.

### 4.7. 3D Cell Culture

All 3D cell cultures were performed in 96 well dishes as described in Murali et al. (2019). In brief, sterile 10X PBS, 1M NaOH and water were pre-warmed in a 37 °C water-bath and an aliquot of 3.2 mg/mL bovine collagen I (Advanced Biomatrix) was brought to room temperature. The 96-well dish was pre-warmed in the incubator. The cells were washed with 1X PBS, trypsinized, harvested and counted to determine cell density. To prepare collagen at a final concentration close to 2 mg/mL from 3.2 mg/mL stock, we combined 100 μL 10X PBS, 10 μL 1 M NaOH, 250 μL water and 640 μL of 3.2 mg/mL collagen stock. The pH was measured to be between 7 and 7.4 using pH strips. The cells were then set up at 30,000 cells per well and incubated for 30 min in 37 °C incubator to allow collagen polymerization. Upon collagen polymerization the media was then added and incubated overnight and drug added the next day.

### 4.8. 10 μM screen of melanoma candidates in 3D

A375 cells were prepared in 3D as described above. For the 10 μM screen, a 1 mM stock drug concentration was first prepared in DMSO. The cells were then incubated at a final concentration of 10 μM with bosutinib, crizotinib, dasatinib, masitinib, ponatinib, sorafenib, pazopanib, nilotinib, sunitinib, SU11652 and a Raf-Mek inhibitor combination of 1 μM dabrafenib + 100 nM trametinib abbreviated as RMIC. The cells were incubated with these drugs for 72h following which viability assays were performed. To determine the rate of proliferation, the cells after 48h of drug treatment EdU at a final concentration of 10 μM was added. This was subsequently incubated in cells for a further 24h to make a total drug incubation of 72h.

### 4.9. Isolating PBMCs from buffy coats

Buffy coats acquired from Carter Blood Care was first diluted 1:4 with 1X PBS. 10mL of the diluted buffy coat was then underlaid with 3mL of Ficoll-Paque PLUS. The Ficoll treated buffy coats were then centrifuged at 2000 rpm for 20 minutes. The PBMCs form a distinct intermediate layer which was carefully removed without disturbing the erythrocytes and resuspended in RPMI with 2% FBS. The sample was then centrifuged at 1500 rpm for 5 minutes to get a PBMC pellet which was then resuspended in RPMI with 10% FBS and interleukin-2 (IL-2) diluted in 20 mM acetic acid at and used at a final concentration of 0.2 μg/mL.

### 4.10. Dose dependent cytotoxicity assay in 3D collagen

To evaluate SU11652 as a therapeutic candidate for melanoma, SU11652 treatment as a function of concentration was performed. A375, M405, M498, M481 melanoma cell lines and HFF1 controls were treated with SU11652. To assess the specificity of CaVu, we used MCF7, S1590 and S159T breast cancer cells since breast cancers were predicted not to be sensitive to SU11652. All cells were treated with SU11652 at 0.01, 0.1, 1, 5, 7.5 and 10 μM along with a DMSO control for 3 days following which imaging based viability assays were performed.

### 4.11. Dose dependent cytotoxicity assay in suspension

PBMCs were cultured in 24-well, ultra-low-adhesion dishes at 100,000 cells/well. The cells were then treated with SU11652 as a function of concentration at 0.01, 0.1, 1, 5, 7.5 and 10 μM along with a DMSO control for 3 days following which imaging based viability assays were performed.

### 4.12. Dose dependent viability assays

To measure the effects of TC-E 5008 on cell viability, breast cancer cells were seeded in 96-well plates (4 x 10^3^ cells/well) in RPMI medium containing 10% fetal bovine serum. After an overnight incubation, cells were treated with varying concentrations of TC-E 5008 or other mutant IDH1 inhibitors for 72 h. Viability was measured using Cell Titer-Glo Luminescent Cell Viability Assay (Promega) in 96-well, flat, clear-bottom, opaque-wall micro plates according to manufacturers protocol.

### 4.13. Statistical tests

All in vitro statistical measurements were performed using ordinary one-way ANOVA, with α = 0.05. For *in vitro* viability assays, the statistical significance was measured by comparing the sum of dead pixels from apoptosis (caspase 3/7) and ethidium homodimer of DMSO treated samples to that of the drug treated samples. For *in vitro* proliferation assays the statistical significance was measured the percent proliferation of DMSO to that of drug treated samples. Results are displayed using *p*-value of < 0.05 : *, < 0.005 : **, and < 0.0005 : ***. We assessed the statistical significance of the difference between mean tumor volume in vehicle-treated versus drug-treated mice bearing M405 and A375 tumors using Student’s unpaired t-test. Results are displayed using *p*-value of \**p* < 0.05, \*\**p* < 0.01, \*\**p* < 0.005.

### 4.14. Evaluating SU11652 in mouse splenocytes

Female C57BL/6 mice (7 weeks old) were purchased from the Jackson Laboratory and were housed in specific pathogen-free conditions at the University of Texas MD Anderson Cancer Center Animal Research Center facilities. Spleens from naive C57BL/6 mice were harvested. Erythrocytes present in the splenocytes were lysed using RBC Lysis Buffer (R7757, Sigma Aldrich). Cells are washed in cRPMI (RPMI+L-glu+Pen/Strep+10%FBS) to remove dead cells. Splenocytes were plated in 1,000,000 cells/mL in cRPMI supplemented with 0.5 μg/mL anti-CD3 (Clone 145-2C11, Biolegend), 0.5 μg/mL anti-CD28 (Clone 37.51, Biolegend), and 100 U/mL rmIL-2 (402-ML-100, R&D Systems) in the presence of increasing concentrations of SU11652 (sc-204310, Santa Cruz Biotechnology) or vehicle control (DMSO) for 72 h. Cells were removed from the plates after the culture period, washed with cRPMI and counted using the Cellometer Auto 2000 Cell Viability Counter (Nexcelon). Viability is assessed by trypan blue exclusion. Experiments in this study were approved by the UT MD Anderson Institutional Animal Care and Use Committee.

### 4.15. 3D imaging based viability and proliferation assays

For viability measurement following drug treatment, media containing drug was aspirated and the cells were incubated with 4 μM EtHd, 2 μM CellEvent Caspase 3/7 and 15 μg/mL Hoechst 33342 in fresh phenol red free DMEM supplemented with 10% FBS. Imaging was then performed with a Nikon Ti epifluorescence microscope with an OKO temperature and CO_2_ control system regulated at 37 °C with 5% CO_2_. The cells in 3D were imaged with a z-step size of 2.5 μm and a total of 201 steps. The filter set for the red channel had an excitation from 540-580 nm and emission at 600-660 nm, green channel had an excitation from 465-495 nm and emission from 515-555 nm and blue channel had an excitation at 340-380 nm and emission at 435-485 nm. To identify cells undergoing S phase of the cell cycle upon drug treatment, we incubated cells with EdU for 24h and labeled using click chemistry. Click reaction was performed as described in (Murali et al., 2019).

### 4.16. Viability imaging of cells in suspension

To evaluate the viability of PBMCs treated with SU11652 as a function of concentration, the cells were incubated with 4 μM EtHd, 2 μM CellEvent Caspase 3/7 and 15 μg/mL Hoechst 33342 for 30 mins. A drop of this cell suspension was then put on a glass slide and placed under a glass coverslip to have the cells in the same imaging plane. Imaging was then performed with the Nikon Ti microscope.

### 4.17. 3D image analysis

We developed custom open source software (https://github.com/Cobanoglu-Lab/FoRTE) to analyze 3D images. In brief, the pipeline corrects for illumination effects and quantifies the pixel-level signal from each z-stack. Specifically, we first correct for the epifluorescence microscope signal artifacts by using a local Gaussian with a large variance to smooth the image. We then subtract this background from the foreground and threshold any resultant negative values to zero. We then find a threshold value for the whole stack. To do this, we first threshold the images at the center of the z-stack to avoid any irregularities that might be present at the end of the stacks. To automatically threshold a single image we use Yen’s algorithm (Yen et al., 1995). For the middle stacks, we compare the threshold value acquired in each image to the mean of the background in the same image. If the threshold is less than ten-fold when compared to background, then we posit that the signal to noise ratio is weak due to the absence of any meaningful biological signal. When we find a robust threshold, we use that across the entire z-stack.

### 4.18. Suspension image analysis using cell profiler

To analyze viable suspension cells upon drug treatment we used a Cell Profiler V2.2 with the pipeline described in Murali et al. (2019). We developed a pipeline that accepts multi-point images to identify for positive pixels in each channel. We used a robust background thresholding method to distinguish foreground (i.e. marked cells) and background regions. The pixel areas of foreground regions were measured for each channel.

### 4.19. Quantifying imaging results from python script and cell profiler

To quantify the results we used total positive pixel counts as the read counts for each channel. We acquired the total hoechst, caspase 3/7 (apoptosis) pixels and EtHd (complete cell death) pixels. To avoid counting cells twice in any channel, we also acquired intersection of pixels between the channels by creating masks between two channels. Overlapping pixels between caspase 3/7 and EtHd were subtracted from caspase 3/7 pixels since cells that are EtHd positive mean they are committed to complete cell death with loss of nuclear membrane integrity. Cells showing overlapping pixels between hoechst and caspase 3/7 and hoechst and EtHd were subtracted from total hoechst pixels since cells entering cells death were distinguished from total hoechst positive cells which then we can consider as total viable pixels. Although there are studies of cells rescued from apoptosis it is not possible to distinguish those cells from the apoptosis committed cells. The raw pixel values for each channel were then quantified by normalizing to the total pixels from the 3 channels of each set to 100%.

### 4.20. Patient-derived and cell line xenograft mouse models

The SU11652 *in vivo* validation using M405 and A375 melanoma cells were performed on NSG mice. Cells were subcutaneously implanted at 3 million cells per mice with matrigel in the lower back of the mice. Once the tumors were palpable (150-200 mm^3^), the mice were randomized and drug treatment was initiated. For the M405 models the mice were treated with vehicle control, 20mg/kg sunitinib and 20mg/kg SU11652. The A375 models were treated with vehicle control, 20mg/kg sunitinib, 20mg/kg SU11652, 50mg/kg dabrafenib + 0.5mg/kg trametinib. Both the M405 and A375 mouse models were treated with drugs daily for the duration of the experiment. Tumor sizes were assessed twice weekly using caliper measurement, and tumor volume was calculated using the formula (length x width2 x π)/6. Mice were euthanized when the tumor volume exceeded 1250 mm^3^ or when necessary for animal welfare. The drug was dissolved in 0.5 % w/v carboxymethylcellulose sodium, 1.8 % w/v NaCl, 0.4 % w/v Tween-80, 0.9 % w/v benzyl alcohol. All mice were sacrificed before tumor weight exceeded 10% of body weight. Animal work described here has been approved and conducted under the oversight of the UT Southwestern Institutional Animal Care and Use Committee.

### 4.21. Syngeneic mouse model studies

Female C57BL/6 mice were purchased from the Jackson Laboratory and were housed in specific pathogen-free conditions at The University of Texas MD Anderson Cancer Center (MDACC) Animal Research Facilities. Animals were allowed to acclimate in the housing facility at least 1 week prior to the experiments. 3×10^5^ B16-F10 tumor cells suspended in 100 μL of phosphate-buffered saline (PBS) were intradermally injected to the flank of the animals. When the tumors were palpable (day 7), tumors were measured and animals were randomized to treatment groups prior to the first treatment dose. SU11652 in DMSO or vehicle control were administered daily via oral gavage at between day 7 and day 10 post tumor inoculation. Tumors were measured by a digital caliper 2 to 3 times per week and the tumor volume was calculated as volume = length x length x width/2. All animal experiments were approved by and performed in accordance with MDACC Institutional Animal Care and Use Committee (IACUC) guidelines.

### 4.22. Patient derived explant studies

Excised tissue samples were processed and cultured *ex vivo* as previously described. Deidentified tumors were obtained from the UT Southwestern Tissue Repository after institutional review board approval (STU-032011-187, 28). Briefly, tumor samples were incubated on gelatin sponges for 24h in culture medium containing 10% FBS, followed by treatment with either vehicle, 10 μM TC-E 5008 for 72h. Representative tissues were fixed in 10% formalin at 4 °C overnight and subsequently processed into paraffin blocks. The sections were then processed for immuno-histochemical analysis.

## 5. Author contributions

V.S. Murali and E.S.W. executed the *in vitro* experiments for the melanoma drug predictions. D.A.C. designed and executed the immunological characterization and syngeneic *in vivo* experiments for SU11652. M.H. and G.V.R. designed the *in vitro* and *ex vivo* validation experiments of the ER+ breast cancer drug predictions. M.Z. performed *in silico* simulations testing the proposed LRTC method. V.S. Malladi and J.G. developed and deployed the CaVu web server. N.S.W. designed and executed the patient-derived explant *in vivo* experiments for SU11652. M.C.C. designed and supervised the study, performed the *in silico* work, and wrote the manuscript.

## 6. Acknowledgements

M.C.C. would like to acknowledge the Lyda Hill departmental start-up funds. D.A.C. is funded as an Odyssey Fellow, supported by the H-E-B Foundation.

## 8. Supplementary figures

**Figure S1:**
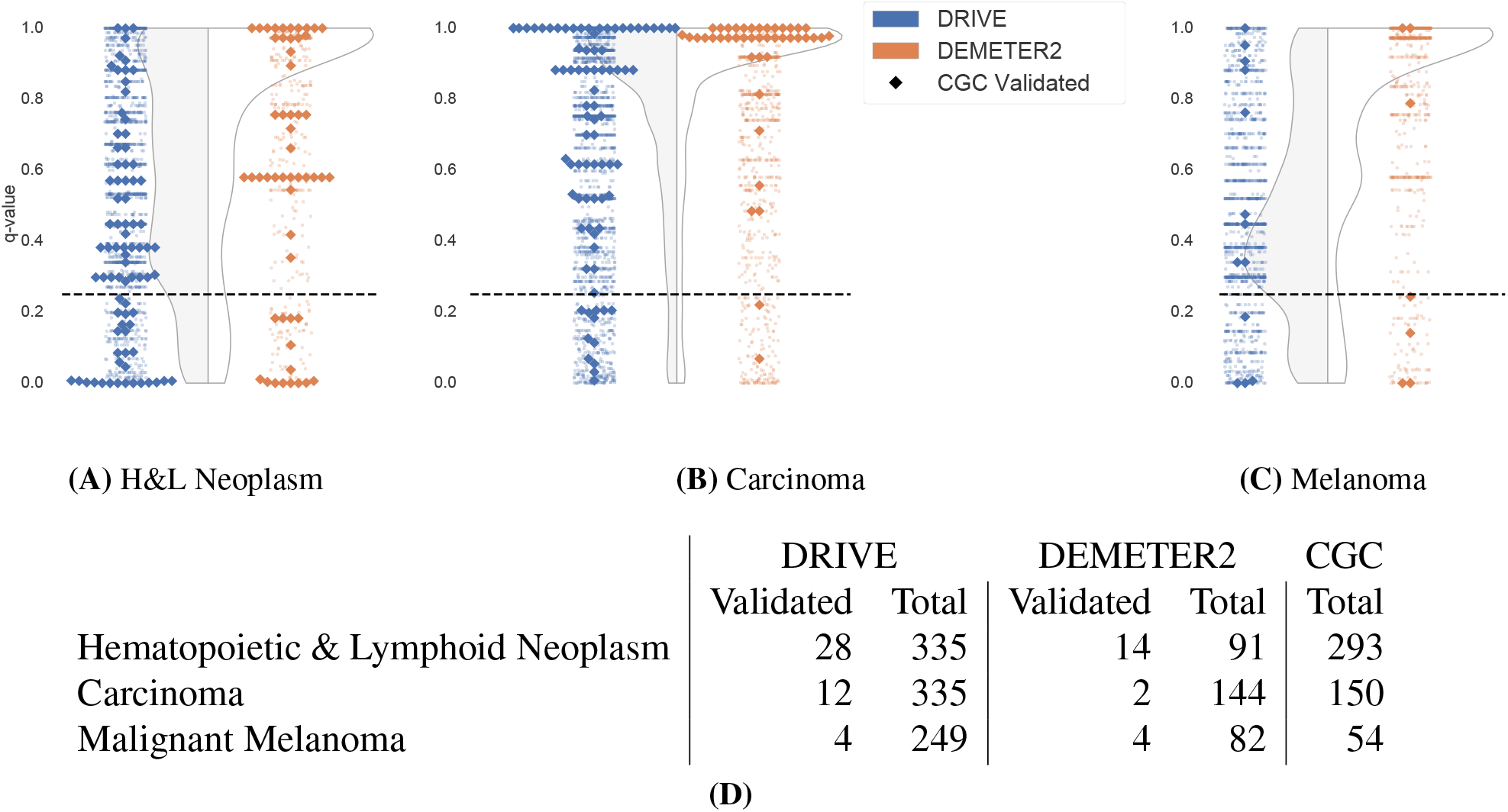
DRIVE is more effective at capturing Cancer Gene Census disease-gene associations than DEMETER2. To compare the two databases for their effectiveness for our purposes of identifying disease-gene associations, we used the same hypergeometric enrichment test we use in our study to identify genes associated with three separate diseases: hematopoietic & lymphoid neoplasm (**A**), carcinoma (**B**), and melanoma (**C**). The FDR adjusted q-values are shown for each gene in each disease context, with a violin plot in the background showing the overall distribution. At q-value cutoff 0.25 (dashed line), the CGC genes for each disease are identified more effectively when using DRIVE as opposed to DEMETER2 (**D**).

**Figure S2:**
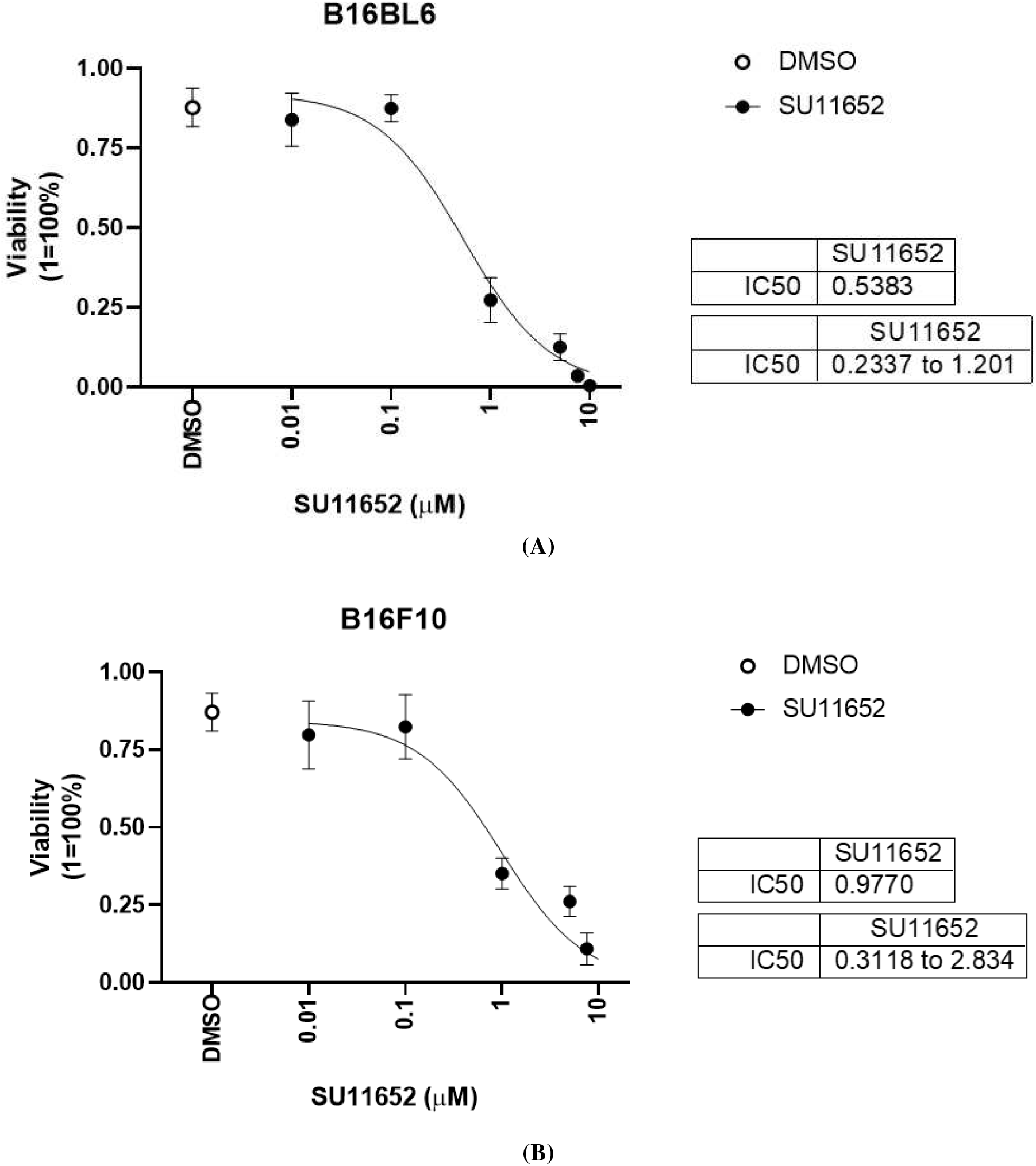
SU11652 is effective on mouse melanoma models B16BL6 (**A**) and B16F10 (**B**).

**Figure S3:**
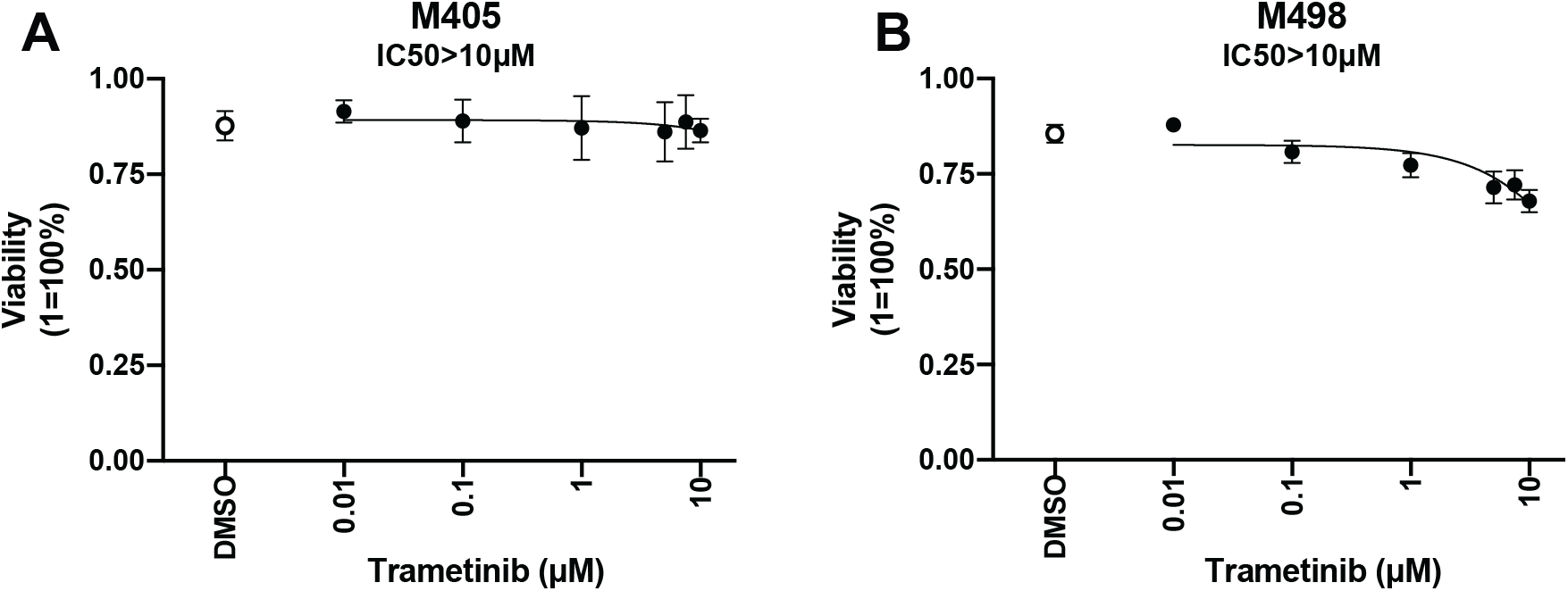
Trametinib dose response on BRAF^WT^ melanoma cell. Dose response viability curves were performed for M405 (**A**) and M498 (**B**) BRAF^WT^ melanoma cells to determine the effect of Trametinib alone along with a DMSO negative control. Statistical significance was measured by comparing the sum of dead pixels from apoptosis (caspase 3/7) and ethidium homodimer of DMSO treated samples to that of the drug treated samples. Results are average ± SEM of three independent repeats with (\**p* < 0.05, \*\**p* < 0.005, \*\*\**p* < 0.0005).

**Figure S4:**
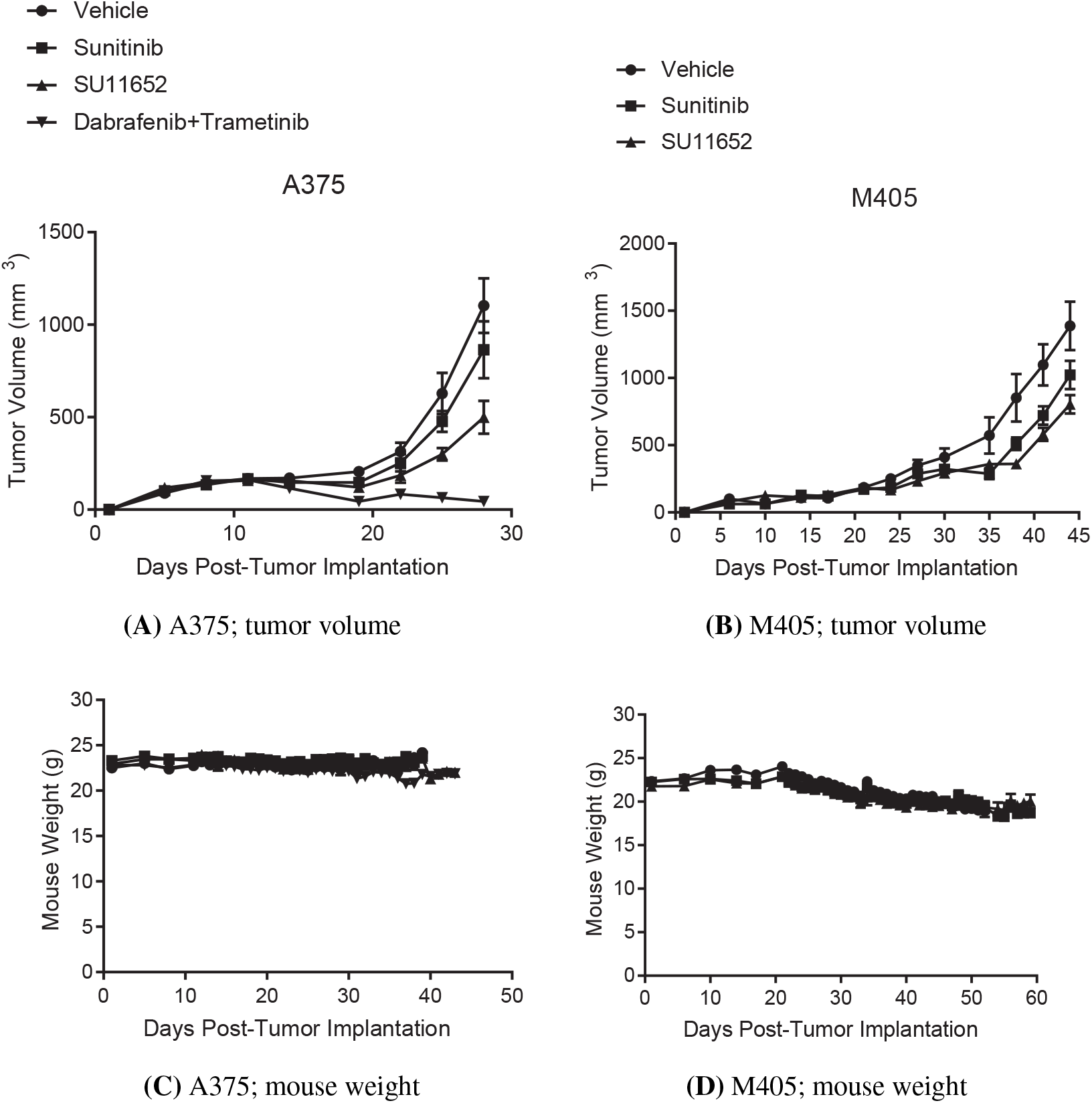
The effect of 20 mg/kg SU11652 treatment on *in vivo* models of human melanoma in immuno-compromised mice on tumor volume and mouse weight. A375 is a BRAF^V600E^ melanoma model cell line, while M405 is a patient-derived xenograft model that is BRAF^WT^ and resistant to existing therapy options.

**Figure S5:**
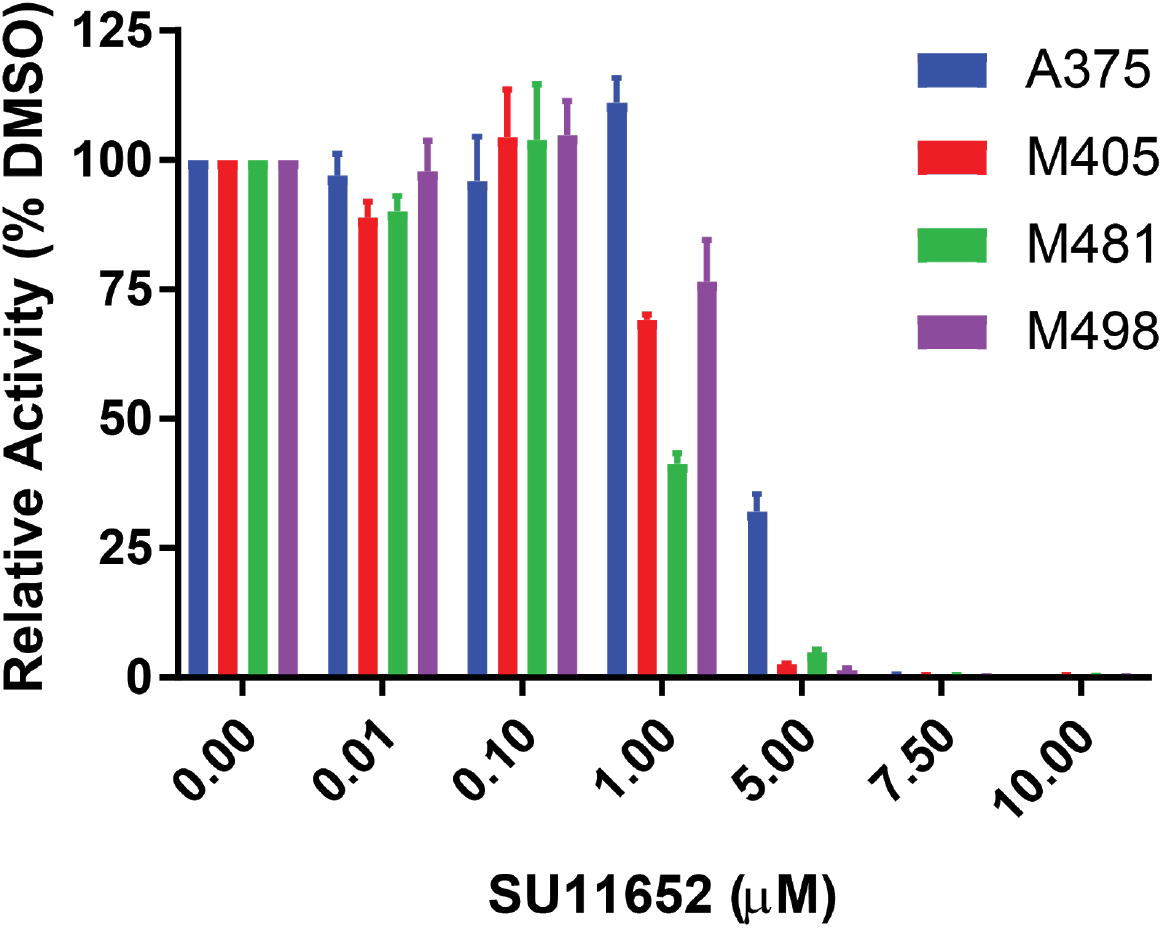
Validating SU11652 with 3D CellTiter-Glo assay. To validate our results with a different and well-established reporter we performed CTG assay. We assessed the impact of SU11652 on four melanoma cell lines: A375, M405, M481 and M498. Plots show average ± SEM of three independent repeats.

**Figure S6:**
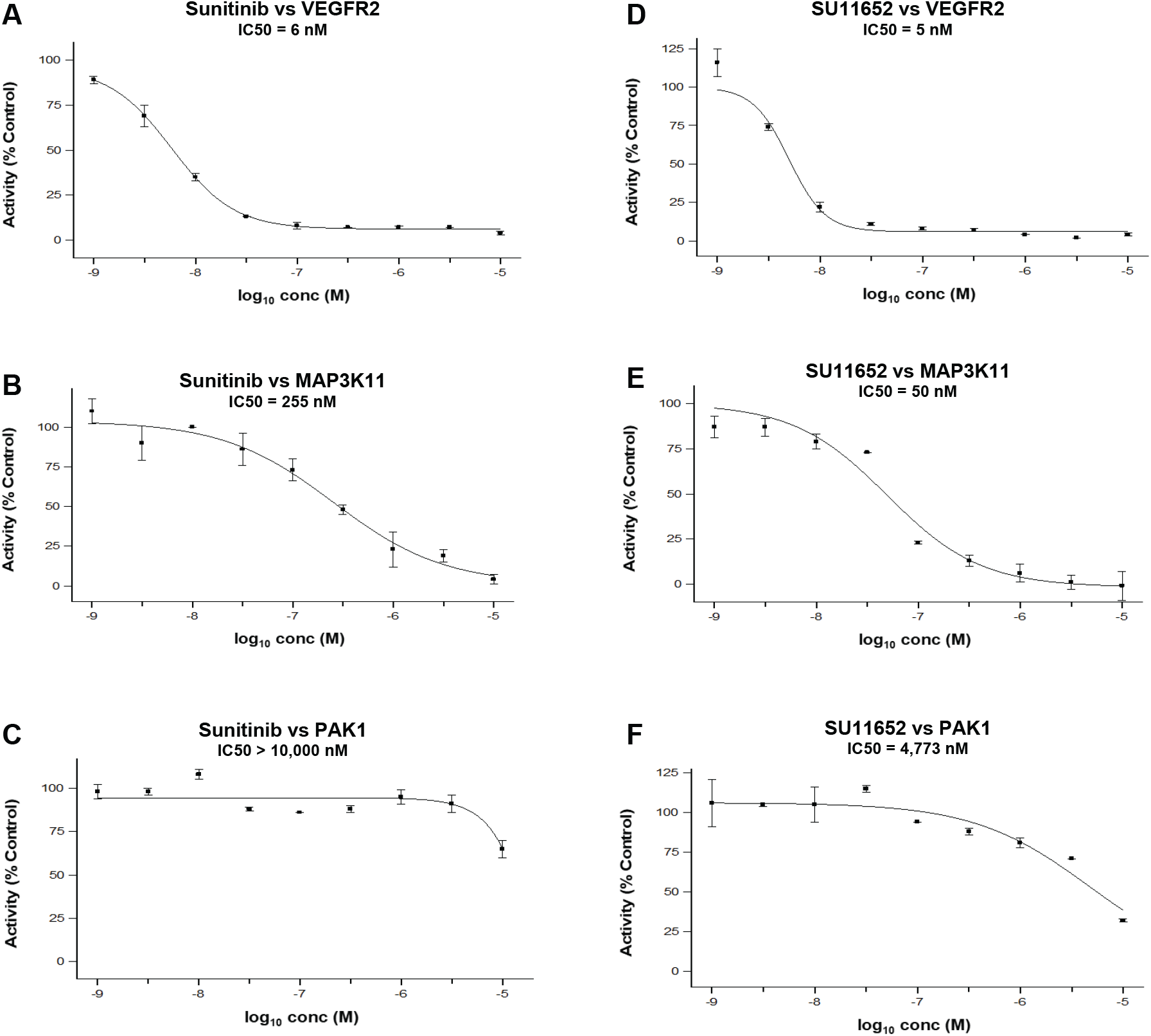
The activity of sunitinib and SU11652 on the three kinases, VEGFR2 (alternatively named KDR), MAP3K11 (alternatively named MLK3), and PAK1. Data collected by EuroFins, through the KinaseProfiler™ and IC_50_Profiler™ services.

**Figure S7:**
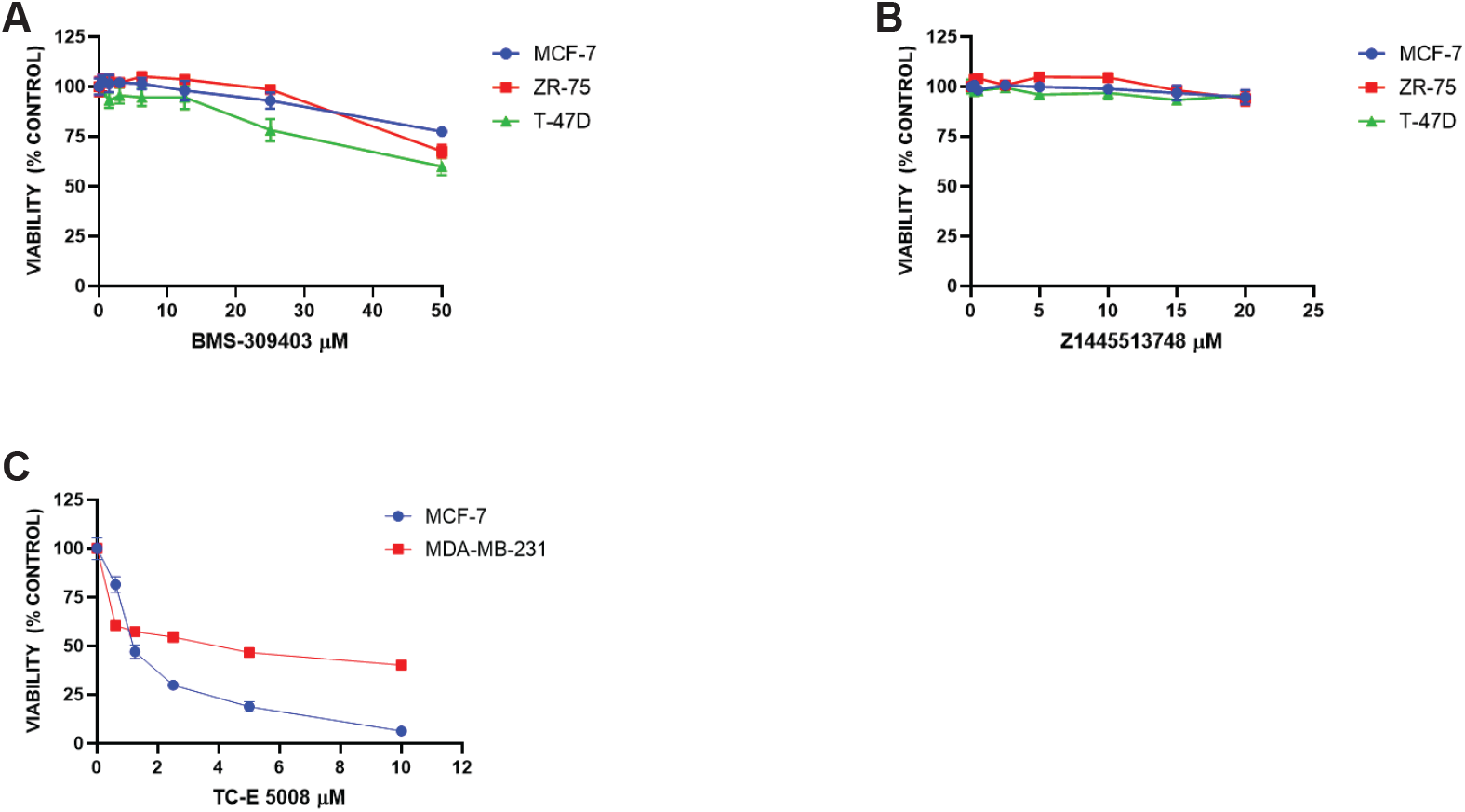
CellTiterGlo assay performed to show anti-proliferative activity of (**A**) FABP5 inhibitor BMS-309403, KDM1A inhibitor Z1445513748 (**B**) on ER positive breast cancer cells and mutant IDH1 inhibitor TC-E 5008 on ER positive (MCF-7) and triple negative (MDA-MB-231) breast cancer cells (**C**).

**Figure S8:**
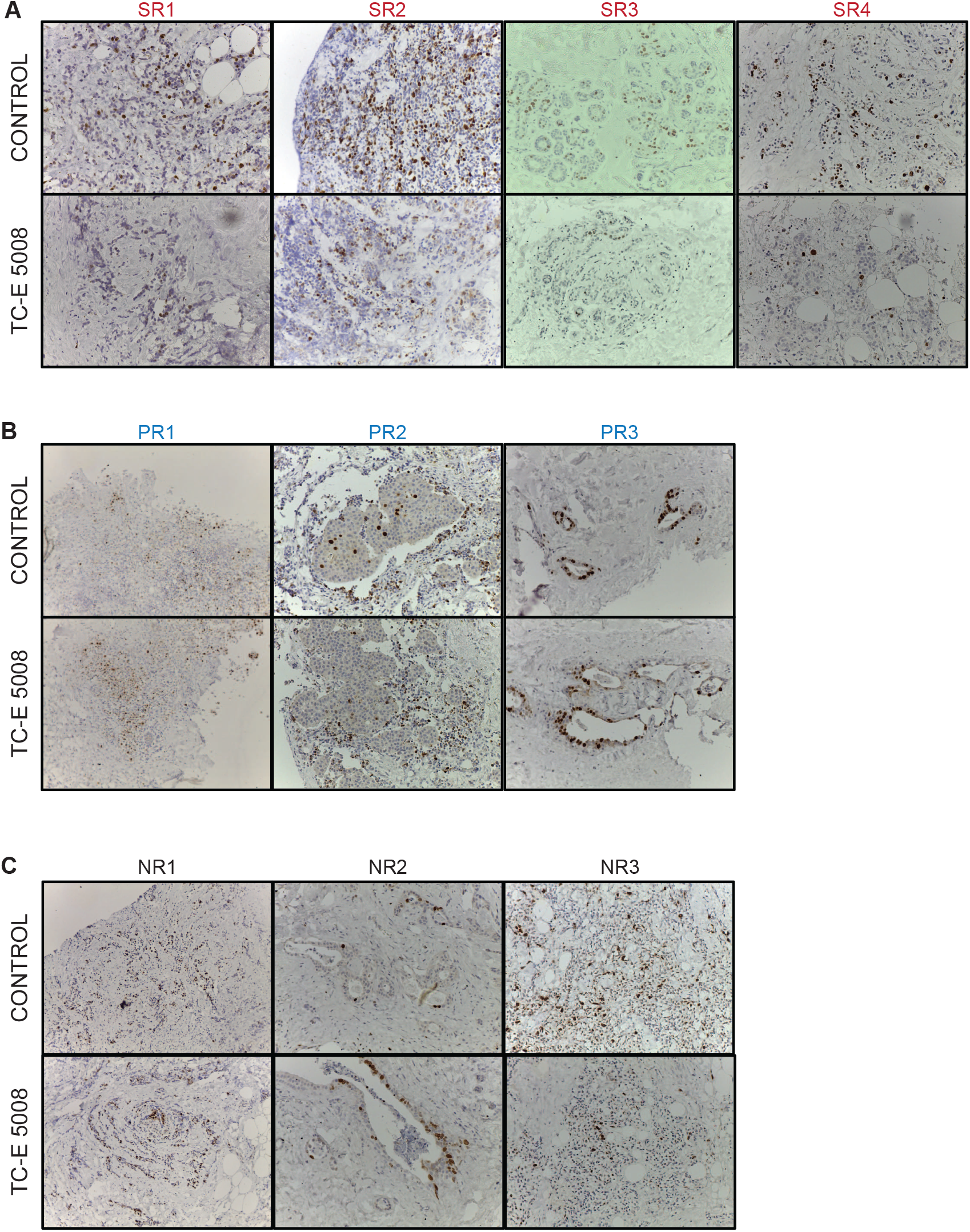
Effect of TC-E 5008 (10 μM) on Ki-67 expression in ER+ tumor explants in (**A**) strong responders (SR), (**B**) partial responders (PR) and (**C**) non-responders (NR). We show one representative from multiple fields of view imaged for each patient & condition.

